# Changes in lipid composition and ultrastructure associated with functional maturation of the cuticle during adult maize leaf development

**DOI:** 10.1101/625343

**Authors:** Richard Bourgault, Susanne Matschi, Miguel Vasquez, Pengfei Qiao, Annika Sonntag, Caleb Charlebois, Marc Mohammadi, Michael J. Scanlon, Laurie G. Smith, Isabel Molina

**Author notes:** **Email addresses** Richard Bourgault; Susanne Matschi; Miguel Vasquez; Pengfei Qiao; Annika Sonntag;, Caleb Charlebois;, Marc Mohammadi; Michael J. Scanlon; Laurie Smith Isabel Molina.

## Abstract

Although extensive prior work has characterized cuticle composition, function, ultrastructure and development in many plant species, much remains to be learned about how these features are interrelated. Moreover, very little is known about the adult maize leaf cuticle in spite of its significance for agronomically important traits in this major crop. We analyzed cuticle composition, ultrastructure, and permeability along the developmental gradient of partially expanded adult maize leaves to probe the relationships between these features. The water barrier property is acquired at the cessation of cell expansion. Wax types and chain lengths accumulate asynchronously along the developmental gradient, while overall wax load does not vary. Cutin begins to accumulate prior to establishment of the water barrier and continues thereafter. Ultrastructurally, pavement cell cuticles consist of an epicuticular layer, a thin cuticle proper that acquires an inner, osmiophilic layer during development, and no cuticular layer. Cuticular waxes of the adult maize leaf are dominated by alkanes and wax esters localized mainly in the epicuticular layer. Establishment of the water barrier coincides with a switch from alkanes to esters as the major wax type, and the emergence of an osmiophilic (likely cutin-rich) layer of the cuticle proper.

**Higlight statement:** Chemical, ultrastructural and functional analysis of cuticle development in partially expanded adult maize leaves revealed important roles for wax esters and an osmiophilic, likely cutin-rich, layer in protection from dehydration.

## Introduction

Plant epidermal cells of the shoot are covered by a hydrophic layer, the cuticle (Esau, 1977), which forms the primary barrier between the plant’s air-exposed surfaces and the external environment. The cuticle limits non-stomatal water loss and gaseous exchanges, protects the plant from extreme temperatures, UV radiation and pathogens, provides mechanical strength, and prevents organ fusion during development (Kolattukudy, 2001; Nawrath, 2002). These functions of the cuticle allowed plants to survive in terrestial habitats early in land plant evolution, with fossil evidence showing cuticles in the earliest known land plants (Bargel *et al.*, 2006).

Plant cuticles contain a lipid biopolymer, cutin, which is embedded in intracuticular waxes and covered by epicuticular waxes (Koch & Ensikat, 2008). Cutin is a polyester matrix composed mainly of glycerol and long-chain (C16 and C18) fatty acid (FA) monomers, usually having ω-linked functional groups and often containing mid-chain hydroxy and epoxy groups. However, the native structure of cutin remains largely hypothetical due to the inability to extract and analyze cutin without prior depolymerization (Kolattukudy, 1980; Graça *et al.*, 2002; Heredia, 2003; Pollard *et al.*, 2008). Cuticular waxes are composed of a mixture of aliphatic and alicyclic compounds with diverse chemistries that are extractable with organic solvents (Samuels *et al.*, 2008; von Wettstein-Knowles, 2012). The aliphatic wax components are usually very-long-chain FAs, alkanes, alcohols, aldehydes, ketones and wax esters. Alicyclic components, including pentacyclic triterpenoids, tocopherols and steroids, are also frequently present (Jetter & Riederer, 2016). The specific composition and quantities of cuticular wax classes may vary significantly across species, plant organs, and developmental stages, and may undergo dynamic effects in response to growth conditions, physical disturbance or damage, and genetic manipulation (Jenks *et al.*, 2001; Jetter *et al.*, 2006; Kosma *et al.*, 2009).

Ultrastructural studies have led to the definition of three principal layers of mature plant cuticles that are observable by transmission electron microscopy (TEM) (Jeffree, 1996; Yeats & Rose, 2013). The basal, cutin-rich “cuticular layer” is recognized ultrastructurally by the presence of fibrils oriented perpendicular to the plane of the cuticle that stain darkly with osmium tetroxide and consist of polysaccharides (Jeffree, 1996; Guzmán *et al.*, 2014; Mazurek *et al.*, 2017). External to the cuticular layer lies the “cuticle proper” containing cutin impregnated with waxes, but considered to be devoid of carbohydrates and lacking the darkly-staining fibrils that characterize the cuticular layer. However, recent studies highlighted the presence of cellulose and pectins in both layers (Guzmán et al., 2014). Epicuticular wax comprises the outermost layer of the cuticle. This layer may exist as a thin film, imparting a shiny or “glossy” appearance, or may include crystals that impart a dull or “glaucous” appearance if sufficiently abundant (Baker, 1982; Koch & Ensikat, 2008). Epicuticular wax is loosely associated with the rest of the cuticle and can be stripped off with adhesives such as gum arabic (Jetter & Schaffer, 2001) or by freezing in glycerol (Ensikat *et al.*, 2000). Analyses of epicuticular wax have shown that its composition is distinct from that of waxes embedded in the cuticle proper (“intracuticular waxes”) (Jetter & Riederer, 2016; Zeisler-Diehl *et al.*, 2018). Understanding how these features of cuticle organization are related to its composition and function is an active area of investigation.

Impermeability to water is a critical feature of the plant cuticle. It is well established that waxes, rather than cutin, confer the majority of the water barrier property of the cuticle (Schönherr, 1976; Kerstiens, 2006; Isaacson *et al.*, 2009; Jetter & Riederer, 2016). Comparisons between closely related species, and between mutant or transgenic vs. wild type individuals, have shown that cuticle thickness has little impact on its function as a water barrier but that wax composition is critical; in particular, alicyclic waxes, including triterpenoids, tocopherols and steroids, increase the permeability of cuticles to water (Vogg *et al.*, 2004; Buschhaus & Jetter, 2012; Jetter & Riederer, 2016). Very few studies have analyzed cuticle development as a way to investigate how compositional and structural features of the cuticle are related to its water barrier properties.

Prior studies investigating the cuticles of maize leaves have mostly focused on juvenile (seedling) leaves of wild-type and *glossy* mutants, which lack the crystalline epicuticular waxes that are normally found on juvenile (but not adult) leaves. Waxes of juvenile maize leaves are dominated by very-long-chain alcohols (63%), with lower proportions of aldehydes (20%), alkanes (1%), and esters (16%) (Bianchi *et al.*, 1978). However, for agronomically important traits related to cuticle function such as drought tolerance and pathogen resistance, the adult leaf cuticle is likely of far greater significance. Adult maize leaf cuticle composition has not been extensively studied, although existing reports indicate that its wax composition differs substantially from that of juvenile leaves, with much higher proportions of wax esters (42%) and alkanes (17%) and smaller proportions of free alcohols (14%) and aldehydes (9%) (Bianchi *et al.*, 1984). The maize adult leaf cutin polyester it is mainly composed of dihydroxyhexadecanoic acid and typical members of the C18 family of cutin acids, including hydroxy and hydroxy-epoxy acids (Espelie & Kolattukudy, 1979).

In this study, we exploit the developmental gradient of partially expanded maize leaves to investigate cuticle ontogeny in the adult maize leaf. We map the acquisition of water barrier properties onto this gradient and relate this functional maturation to changes in chemical composition and ultrastructure, yielding new insights into composition/structure/function relationships.

## Materials and Methods

### Plant material and growth conditions

For cuticular lipid analysis, B73 maize (*Zea mays*) plants were grown in a 25°C day, 20°C night, in 60% relative humidity with a, 16 h: 8 h, light: dark photoperiod in controlled growth chambers until harvest at approximately 4 wk. For functional maturation and transmission electron microscopy analysis, plants were grown in a glasshouse without supplemental lighting with temperatures in the range of 18-30°C. In preliminary experiments, wax and cutin profiles were found to be very similar at all developmental stages for plants grown under these different conditions, permitting comparisons of results. All experiments presented focused on partially expanded leaf 8 (counting the first seedling leaf as leaf 1) when it was 50-60-cm long with a < 1-cm long (unexpanded) sheath.

### Cell length measurements

Leaf 8 was cut into 2-cm segments and stained with 10 µg ml^−1^ propidium iodide. Pavement cell outlines were visualized by confocal microscopy and their surface areas were measured with Image J. Three fields of view were analyzed at each leaf per stage for each of three individual leaves. The number of individual cells measured along the gradient were: 1 cm = 393, 2 cm = 391, 3 cm = 327, 4 cm = 222, 5 cm = 372, 6 cm = 296, 7 cm = 325, 8 cm = 246, 9 cm = 233, 10 cm = 255, 11 cm = 252, 12 cm = 206, 30 cm = 159, 40 cm = 132.

### Cuticle permeability experiments

B73 leaf 8 was excised, the cut end sealed with petroleum jelly, and submerged in 0.05% TBO in water. After 1 or 2 h, leaves were photographed and successive 1-cm segments were homogenized and extracted with isopropanol/formic acid. To separate TBO from chlorophyll, extracts were treated with hexane. TBO in the aqueous phase was quantified by spectrophotometry (A630 nm). To measure resistance to dehydration, intact plants were kept in the dark for two h to close stomata. To measure water loss, 2-cm segments were photographed and weighed repeatedly over a 2 h period. Weight loss was normalized against surface areas determined with Image J (Schneider *et al*., 2012).

### Transmission Electron Microscopy (TEM)

Leaf tissue pieces (~2 mm × 10 mm) were cut from the area midway between the midrib and leaf margin within each developmental interval analyzed, and fixed in 2% paraformaldehyde-2% glutaraldehyde in 0.1 M sodium cacodylate buffer (pH 7.3-7.5) for 3 d at 4°C. Samples were washed with 0.1 M sodium cacodylate buffer (x4) and postfixed in 1:1 solution of 2% osmium tetroxide in 0.1 M sodium cacodylate for at least 12 h at 4°C. Samples washed with 0.1 M sodium cacodylate buffer (x2) and then water (x2) before second postfixation in aqueous 2% uranium acetate overnight (O/N) at 4°C. Samples washed with water (x2) before acetone dehydration series (20, 40, 60, 80, and 100% 30 min each) on ice and final 100% dry acetone at room temperature. Samples were infiltrated in a acetone:epoxy resin (Durcupan ACM, Sigma) series v/v (3:1 2 h, 1:1 O/N, 1:3 2 h, 0:1 O/N) and embedded in resin at 60°C for at least 3 d. Seventy-nm sections were cut using a Leica UCT ultramicrotome and collected on formvar/carbon copper 100-mesh grids (Electron Microscopy Sciences) with glow discharge and post stained with aqueous 2% uranium acetate and Reynold’s lead citrate 5 min each. Grids were viewed on a JEOL JEM-1400Plus (80kV) TEM equipped with a Gatan OneView camera or with a Tecnai G2 Spirit BioTWIN (80kV) TEM equipped with an Eagle 4k HS digital camera (FEI, Hilsboro, OR).

### Cuticular lipid analysis

#### Cuticular wax extraction

##### Wax extraction method optimization

Wax extraction conditions were first optimized for solvent polarity and extraction time (Loneman *et al.*, 2017). A mature portion of the leaf (18-24 cm) was used for all extractions. Three solvent systems were compared: hexane:diethyl ether (9:1) [2.12], chloroform:hexane (1:1) [3.35], and pure chloroform [4.81], listed in the order of increasing polarity (dielectric constants shown in square brackets). Leaf segments were immersed in each solvent and gently shaken for 1 min. To further optimize the immersion time, tissues were extracted with chloroform for 0.5, 1, 2 and 5 min. Because no significant differences were observed with any of the solvents or extraction times analyzed (Fig. S1), 1 min dipping in chloroform was used for all subsequent wax determinations.

##### Developing leaf wax extraction

Two-cm-long pieces (excised between 2 and 22 cm from the leaf base) were immersed in 5 ml chloroform and 5 μg each internal standard, namely *n*-tetracosane (24:0 alkane), 1-pentadecanol (15:0-OH) and heptadecanoic acid (17:0), were added to each extract. The chloroform extracts were then evaporated under a gentle stream of nitrogen. Each leaf portion was scanned and its surface area measured using ImageJ and multiplied by two to account for both surfaces.

### Isolation of epicuticular and intracuticular waxes

Epicuticular waxes were removed by applying a gum arabic solution on the surface (Jetter and Schäffer, 2001). Intracuticular waxes were calculated by subtracting epicuticular waxes from total chloroform-extracted cuticular waxes. Gum arabic (Acros Organics) was washed with methylene chloride using a Soxhlet extractor at 40°C for 48 h subsequently used to prepare a 0.6 g ml^−1^ solution in sterile deionized water. Twelve plants were employed to produce 4 biological replicates per sample. After removing the mid-rib, one half of each leaf segment was used to extract epicuticular waxes and the other half was immersed in chloroform for total cuticular wax analysis. Gum arabic solution was applied to four 3-cm-long segments corresponding to 4-7, 8-11, 12-15 and 19-22 cm from the leaf base; for each sample, the gum was applied to two pieces on the adaxial side and two pieces on the abaxial side. Gum pieces were peeled off and photographed alongside a ruler to determine surface area using ImageJ software. Gum pieces were extracted with chloroform and waxes analyzed following the same procedure described above.

### GC-FID and GC-MS analysis

Wax extracts were transformed into their trimethylsilyl (TMS) ester and ether derivatives following the protocol described by Razeq et al. (2014). For chemical identifications, the samples were analyzed by gas chromatography-mass spectrometry (GC-MS) on a TRACE 1300 Thermo Scientific gas chromatograph with a Thermo Scientific ISQ Single Quadrupole mass spectrometer detector. Splitless injection was used with a VT-5HT capillary column (30 m × 0.25 mm i.d., and 0.10 µm film thickness) and a helium flow set at 1.5 ml min^−1^. Temperature settings were as follows: inlet 330°C, detector 300°C, oven temperature set at 150°C for 3 min and then increased to 330°C at a rate of 3°C min^−1^, then from 330°C to 390°C at 6°C min^−1^, with a final hold at 390°C for 10 min. For compound quantification, wax samples were analyzed on a Thermo Scientific TRACE 1300 gas chromatograph coupled to a flame ionization detector (GC-FID) using similar column and chromatographic conditions to those described for the GC-MS analysis. Wax components were identified by their relative retention times and characteristic mass spectra by comparison to published MS data (e.g., Christie, 2019) or by searching a mass spectral reference library (NIST 2011). The approach used to determine the double bond position of the alkenes identified in our analysis is described in Methods S1. Wax components were quantified based on the total FID ion current. Theoretical correction factors were applied that assume that the FID response is proportional to carbon mass for all carbons bonded to at least one H-atom (Christie, 1991). Because split/splitless injection is known to bias against high molecular weight compounds, calibration curves for wax esters (WEs) with even-chain standards were generated and used to calculate the amounts of WEs found in the samples (Method S2; Fig.S2).

### Lipid polyester analysis

Leaf segments used for wax extractions were delipidated by shaking in isopropanol and chloroform:methanol mixtures, as described in Jenkin and Molina (2015). Polyester monomers released by transesterification of solvent-extracted tissues were transformed into silylated derivatives, and analyzed by GC-MS (Jenkin & Molina, 2015). Cutin monomers were identified by their relative retention times and characteristic fragmentation patterns, and by comparison of the EI-MS spectra of their TMSi derivatives with published spectra (Eglington and Hunneman, 1968; Eglinton *et al.*, 1968; Holloway and Deas, 1971; Holloway and Deas, 1973; Deas *et al.*, 1974). Control experiments to assess the monomer contribution by other leaf tissues (i.e., bundle sheath suberin) are detailed in Methods S3.

## Results

### The adult maize leaf cuticle acquires its water barrier property at the cessation of cell expansion

The maize leaf develops from tip to base in a continuous gradient, providing an excellent system for comparative analysis of different developmental stages in a single leaf at one time point (Sylvester & Smith, 2009). For this study, we chose leaf 8, the first fully adult leaf, in a partially expanded state (50-60cm long) of the inbred line B73 as the standard for all experiments. Cells are still dividing at the base of this leaf, with later developmental stages (cell expansion, differentiation, maturation) represented at successively more distal positions (Facette *et al.*, 2013). Two approaches were taken to investigate the establishment of water barrier properties of the cuticle along this gradient (Fig. 1). In the first approach, intact leaves were stained with the water-soluble dye toluidine blue-O (TBO) (Tanaka *et al.*, 2004). TBO penetrated the developing cuticle until ~10-12 cm above the leaf base (Fig. 1a, b). In a second approach, dehydration rate in the dark (where stomata are closed, if present, so that water is lost mainly by transpiration across the cuticle; Ristic & Jenks, 2002), was measured for successive 2-cm intervals and was also found to decrease gradually from the leaf base to ~10-12 cm (Fig. 1c). To relate these findings to cell expansion, we measured pavement cell areas along the developmental gradient. Almost all cell elongation occurred between 3 and 12 cm from the leaf base with the highest cell expansion rate between 4 and 8 cm (Fig. 1d) where there is a local increase in TBO permeability (Fig. 1a, b). Together, these results showed that the water barrier property of the adult maize leaf cuticle is fully established ~10 cm above the leaf base, coinciding with the completion of cell elongation and shortly before emergence of the leaf from the whorl (point of emergence, or POE at 17-18 cm from leaf base; Fig. 1a).

**Figure 1.**
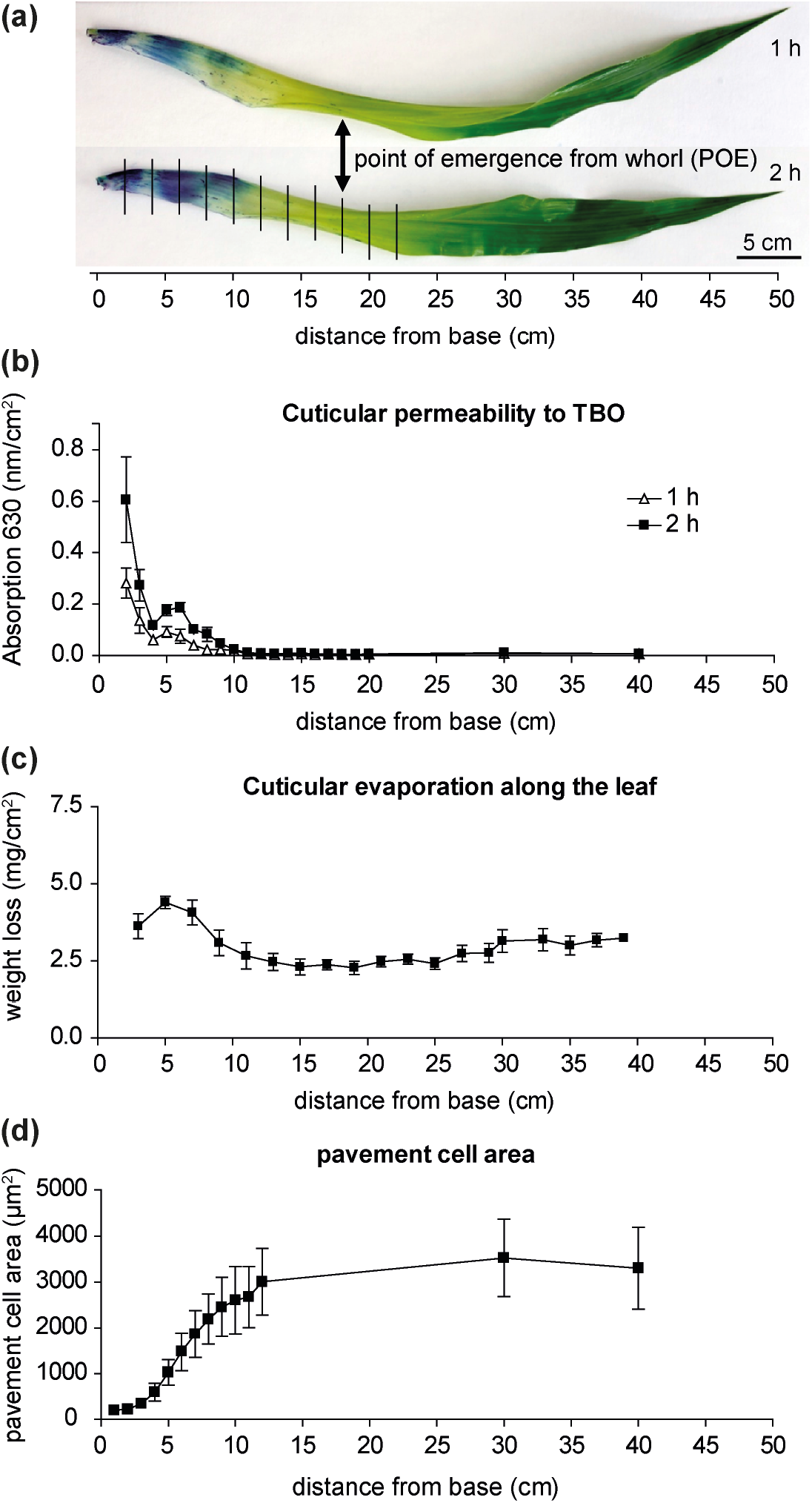
Cuticle maturation along the developmental gradient of a partially expanded, adult maize leaf. (a) Partially expanded, intact adult maize leaves (B73, leaf 8 at 50-60 cm length) stained with TBO for 1 or 2 hours. Vertical lines indicate segments harvested for subsequent analysis of cuticle composition and ultrastructure. (b) Quantification of TBO penetration of the cuticle along the leaf developmental gradient (n = 3 leaves per time point). (c) Cuticular evaporation rate, a measure of dehydration resistance, was measured for successive 2 cm segments of the leaf developmental gradient (n = 3 leaves per time point, mean +/-SE). (d) Pavement cell surface areas along the leaf developmental gradient. Cell outlines were measured with Image J in confocal microscopy images of propidium iodide-stained leaf segments. Three fields of view were analyzed for each of 3 individual leaves (n > 130 cells measured at each position analyzed).

### The timing of accumulation of different wax types and chain lengths varies widely along the adult maize leaf developmental gradient

To elucidate the compositional changes associated with cuticle ontogeny in adult maize leaves, we analyzed both cuticular waxes and cutin along this developmental gradient. Cuticular waxes are, by definition, organic-solvent-soluble, while cutin is insoluble (Kolattukudy, 1980). Most plant surface waxes are efficiently extracted by quick immersion in chloroform (Holloway, 1984; Riederer & Schneider, 1990; Jetter *et al.*, 2006; Fernandez-Moreno *et al.*, 2016; Hegebarth & Jetter, 2017). However, it is unclear if other solvents or combinations of solvents that are more or less polar than chloroform can more efficiently extract the main wax components identified in maize mature leaf samples, namely hydrocarbons, wax esters, fatty acids, aldehydes and fatty alcohols (Bianchi *et al.*, 1984). Following comparisons of extraction methods to optimize wax yields (Fig. S1), waxes were extracted from successive 2-cm intervals corresponding to the segment between 2 and 22 cm from the leaf base.

Given the immaturity of the cuticle between 2 and 4 cm from the base of the emerging leaf 8, wax loads at this position were surprisingly high (109 ± 6 μg dm^−2^). Overall wax load did not increase across the developmental gradient with amounts fluctuating between 94 and 115 μg dm^−2^ (Fig. 2). However, wax composition varied dramatically across the gradient. Hydrocarbon (alkane/alkene) coverage was highest 2-4 cm from the leaf base, dropping by 70% from there to 10-12 cm (Fig. 2; Table S1). Alkyl esters followed the opposite trend, accumulating from nearly undetectable levels initially to become the most prevalent class of waxes by the 10-12 cm stage when the water barrier property is fully established, reaching a coverage of about 59 μg/dm^2^ at that position, and remaining at this level thereafter. Fatty alcohols reached their highest concentration between 6 and 8 cm from the leaf base (14.2 μg dm^−2^) and subsequently decreased, possibly because alcohols are incorporated into wax esters. Free fatty acids, aldehydes, and alicyclics represented a small fraction of waxes at all stages and did not change greatly in abundance across the developmental timecourse analyzed (Fig. 2; Table S1).

**Figure 2.**
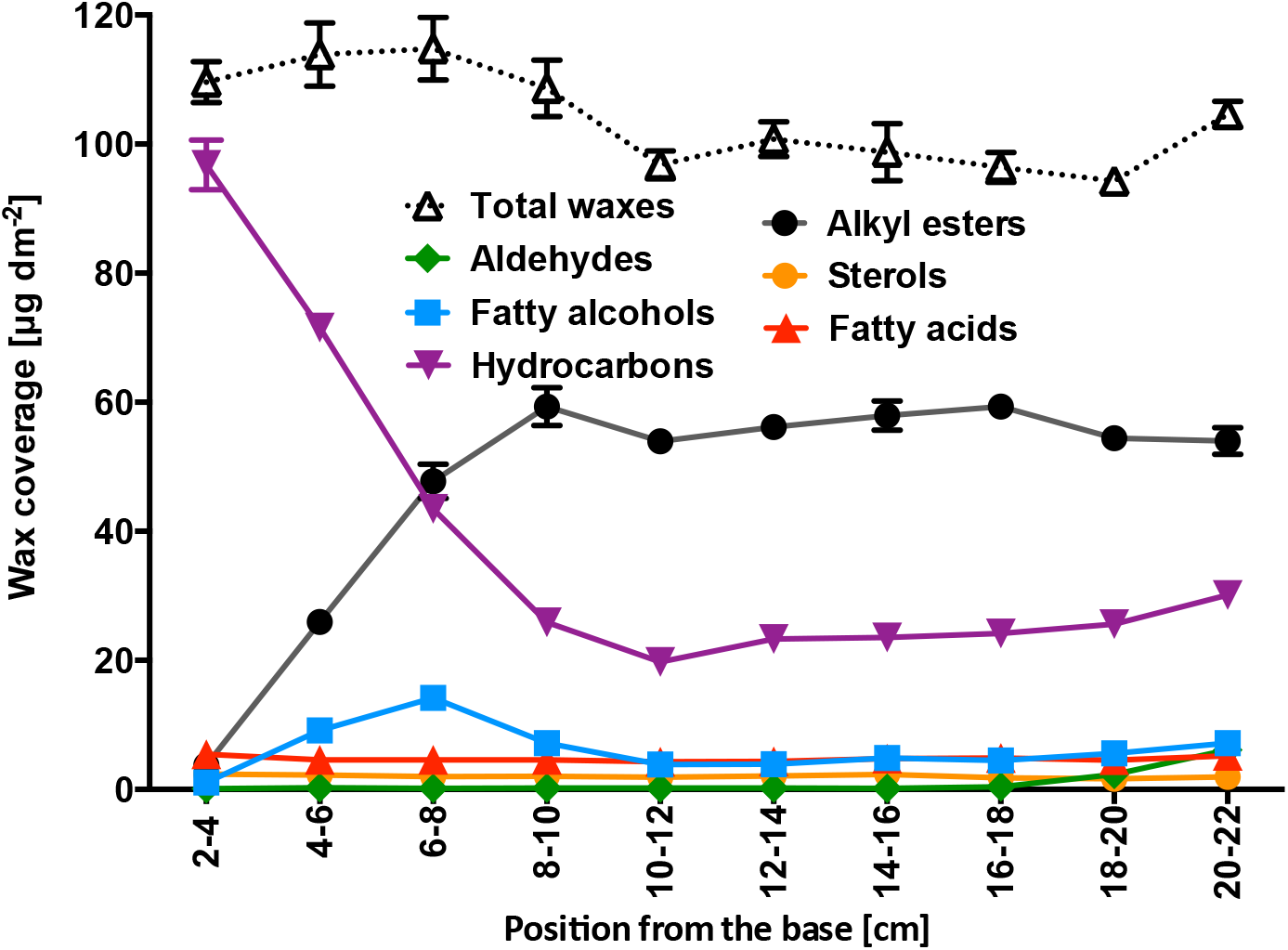
Cuticular wax composition along the adult maize leaf developmental gradient. Changes in the accumulation of six different classes of compounds present in chloroform-extracted wax mixtures. Mean of 4 replicates and SE are reported.

Conspicuous compositional changes were observed within the hydrocarbon series. C_21:0_, C_23:0_ and C_25:0_ alkanes, as well as C_29:1_ and C_31:1_ alkenes, were the predominant hydrocarbons at the earliest stages analyzed. However, these components decreased in abundance gradually across the gradient, reaching low abundance by the 10-12 cm stage when the water barrier property is fully established, and nearly disappearing thereafter (Fig. 3a; Table S1). Each C_29:0_ and C_31:0_ alkene included two positional isomers –determined by mass spectrometry analysis of their corresponding dimethyl disulfide adducts (Buser *et al.*, 1983)– namely 9- and 10-nonacosene and 9- and 10-hentriacontene (Fig. S3). C_27:0_ and C_29:0_ alkanes also declined in abundance from 2 to 12 cm remaining constant thereafter. The longest alkanes (C_31:0_ to C_37:0_) were almost undetectable until the 8-10 cm stage, and also increased in abundance thereafter. Thus, we observed an overall shift from shorter to longer hydrocarbons as development proceeded, with chain lengths <30C predominating initially but shifting to >30C by the time cuticle development was complete.

**Figure 3.**
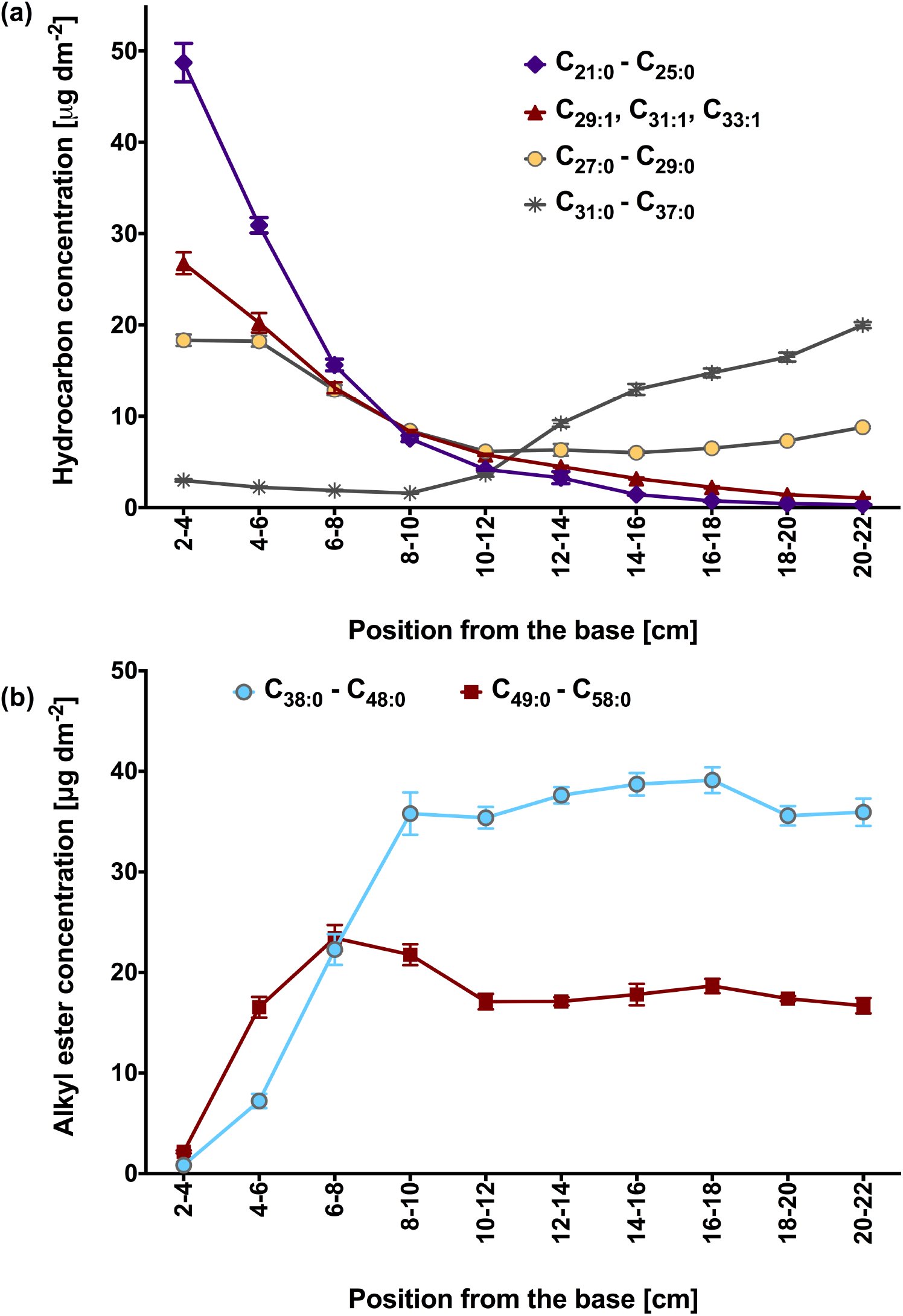
Hydrocarbon and alkyl ester composition along the adult maize leaf developmental gradient. (a) Hydrocarbon (alkane and alkene) composition. (b) Alkyl ester composition (low abundance components, namely C38 and odd-carbon number esters, are not shown). Mean of 4 replicates and SE are reported.

Although alkanes were the dominant class 2-8 cm from the base, alkyl esters were the most prevalent group of waxes beyond that point (Fig. 2; Table S1). Two groups of wax esters were found, one containing the C_38_ to C_48_ homologues (including small proportions of odd-carbon numbered species), and a second composed mostly of C_50_ to C_58_ homologues. The distribution and amount of these two groups differed along the gradient (Fig. 3b). In contrast to findings for alkanes, the longer alkyl esters accumulated first, reaching their peak abundance by 6-8 cm and declining slightly thereafter, while the shorter C_38_-C_48_ group of alkyl esters reached nearly peak levels by the 8-10 cm stage with little or no further changes thereafter (Fig. 3b; Table S1). Furthermore, this shorter chain WE group accumulated to ~twice the abundance of the longer chain group (Fig. 3b). Each alkyl ester class was composed of a mixture of isomers. Overall, the predominant acid and alcohol moieties at 20-22 cm from the leaf base were 18:0- 22:0 (92 % of the esterified fatty acids) and 22:0-34:0 (86% of the esterified alcohols), respectively (Table S2).

The composition of the free fatty alcohol fraction (i.e., primary alcohols not incorporated into alkyl esters) reflected that of the alcohols that are incorporated into alkyl esters (Table S1; Table S2); only 22:0 1-alcohol was not found in its free form. In spite of acyl chains varying between 16 and 30 carbons in length in the alkyl esters, only 16:0 and 18:0 free fatty acids were detected in the wax mixtures of any stage analyzed (Table S1). Collectively, the broad chain-length distribution of wax ester homologs and their isomers and the fact that the bulk of primary alcohols and fatty acids are incorporated into wax esters suggests that one or more wax ester synthases efficiently incorporate substrates of various chain-lengths.

The aldehyde fraction comprised an even-numbered homologous series ranging from 26 to 34 carbons in length (Table S1). Unlike the other wax classes, aldehydes only accumulated significantly >18 cm from the base, with an overall amount of only 5.6 % of the total waxes in the segment comprised between 6 and 22 cm from the leaf base.

Alicyclic triterpene derivatives identified in the analyzed leaf 8 developmental stages were largely comprised of sterols, including β-sitosterol, campesterol and stigmasterol (Table S1). Neither the overall amount, nor the relative proportions, of sterols varied substantially across the developmental timecourse analyzed (Fig. S4). Thus, variation in alicyclic wax content does not appear to play a role in establishing the water barrier property of cuticle during adult maize leaf development.

### Cutin and wax deposition along the maize leaf developmental gradient are not synchronized

Unlike waxes, cutin steadily increased between 2 and 18 cm from the leaf base, reaching a surface mass per unit area twice that of the combined waxes and remaining constant thereafter (Figure 4a). Another notable difference compared to wax biogenesis was that the relative proportion of the main two classes of cutin monomers identified, ω-hydroxy acids and α,ω-dicarboxylic acids, remained constant at about 2:1 as both accumulated (Fig. 4a). Largely consistent with an earlier study on of cutin composition in adult maize leaves of a different inbred (Espelie & Kolattukudy, 1979), the major cutin monomers in adult B73 leaves were identified as 9,16-dihydroxyhexadecanoic acid, 18-hydroxyoctadecenoic, 10(9),18-dihydroxyoctadecenoic acid, 9-epoxy-18-hydroxy stearic acid and 9-hydroxy-1,18-octadecene dioic acid, which together accounted for 86 mole % of the cutin load at maturity (Fig. 4b; Table S3). Two of these monomers, 18:1 10(9),18-dihydroxyoctadecenoic acid and 9-hydroxy-1,18-octadecene dioic acid, have been found previously in the depolymerization products of maize leaf cutin (Holloway, 1983), However, these monomers are unlikely to be products of enzyme-catalyzed reactions; instead, they probably result from photo-oxidation and auto-oxidation of unsaturated fatty acids (Kosma *et al.*, 2015). Although aromatics (i.e. ferulic and coumaric acids) were abundant components of the cutin profiles in our analysis utilizing whole leaves, analysis of cutin monomers in enzymatically isolated epidermal tissues indicated that the bulk of these hydroxycinnamic acid derivatives, along with hexadecanoic acid, are of non-epidermal origin (Fig. S5). Therefore, the cuticle composition results reported here do not include ferulate, coumarate and hexadecanoic acid (Table S3).

**Figure 4.**
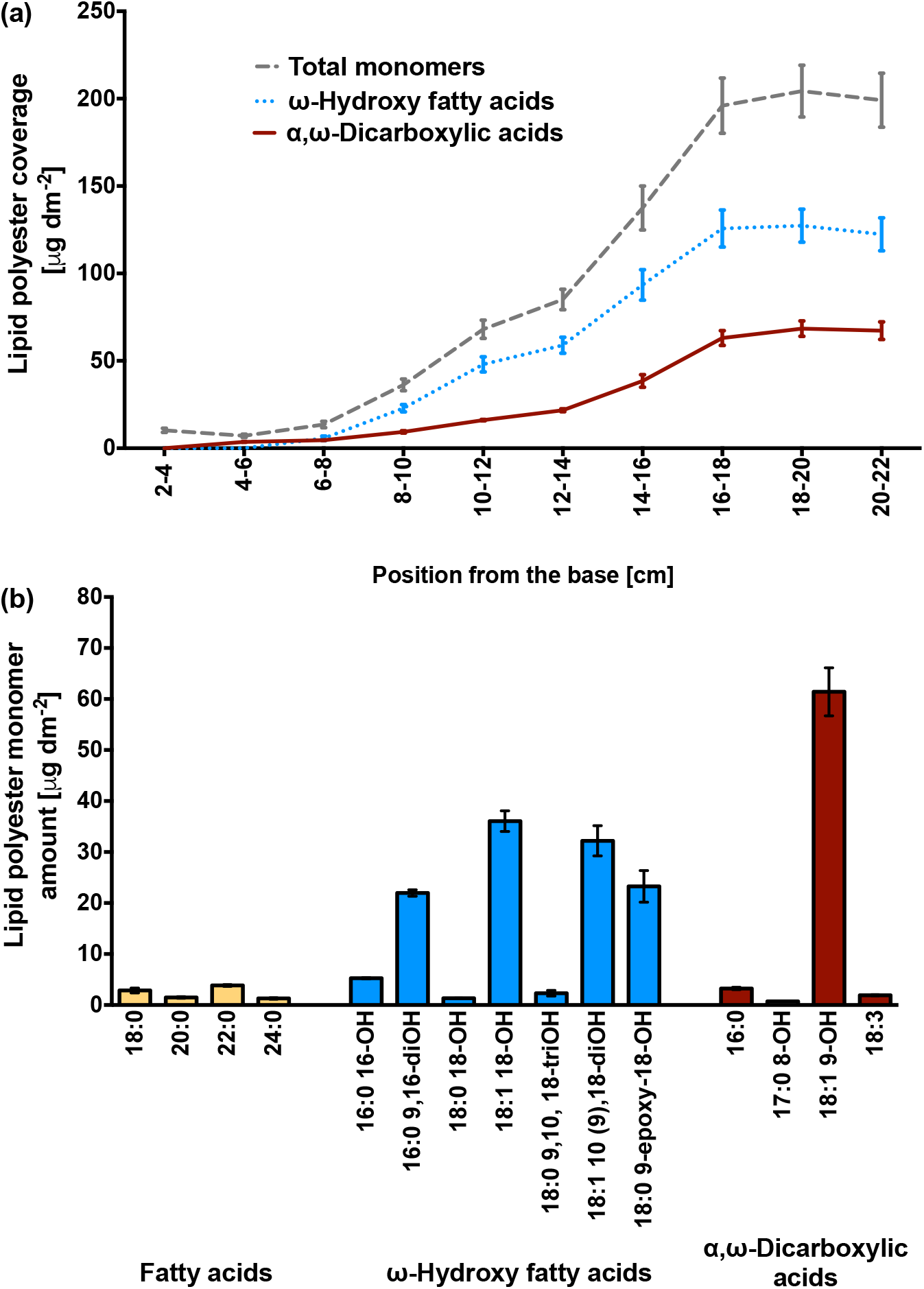
Lipid polyester monomer accumulation along the adult maize leaf developmental gradient. (a) Changes in the accumulation of the two main classes of cutin monomers present in maize leaf cutin compared to the total amount of monomers. (b) Representative profile of the maize mature leaf cutin monomer composition (portion between 20 and 22 cm from the leaf base). Means of 4 replicates and SE are reported.

### Ultrastructural analysis of pavement cell cuticles in the adult maize leaf reveals distinct layers but no cuticular layer

Prior studies of maize leaf cuticle ultrastructure have examined only juvenile leaves, and the published images and reports do not reveal many ultrastructural details (Holloway, 1982; Ristic & Jenks, 2002; Sturaro *et al*., 2005). Ultrastructural images of adult maize leaf pavement cells, revealed a thin cuticle (approximately 40 nm) composed of four zones with distinct osmium staining characteristics (Fig. 5a). The boundary between the cell wall and cuticle is demarcated by a darkly-stained interface (white arrowhead, Fig. 5a), which presumably corresponds to the pectin-rich wall/cuticle interface described for many other plant species (Jeffree, 2006). Another darkly-stained layer (black arrowhead, Fig. 5a) was observed at the outer surface of the adult maize leaf cuticle. This layer was loosely associated with the rest of the cuticle and is removable with gum arabic (Fig. 6a, b), identifying it as epicuticular wax. Between the wall/cuticle interface and the epicuticular layer, darker (asterisk) and lighter-staining zones were observed (Fig. 5a). The lack of fibrils indicate that the darker zone should be classified as a layer of the cuticle proper rather than as a cuticular layer. However, cuticles of specialized cell types (bulliform cells, stomatal guard and subsidiary cells) in the adult maize leaf epidermis are much thicker and exhibit a cuticular layer (Fig. S6).

**Figure 5.**
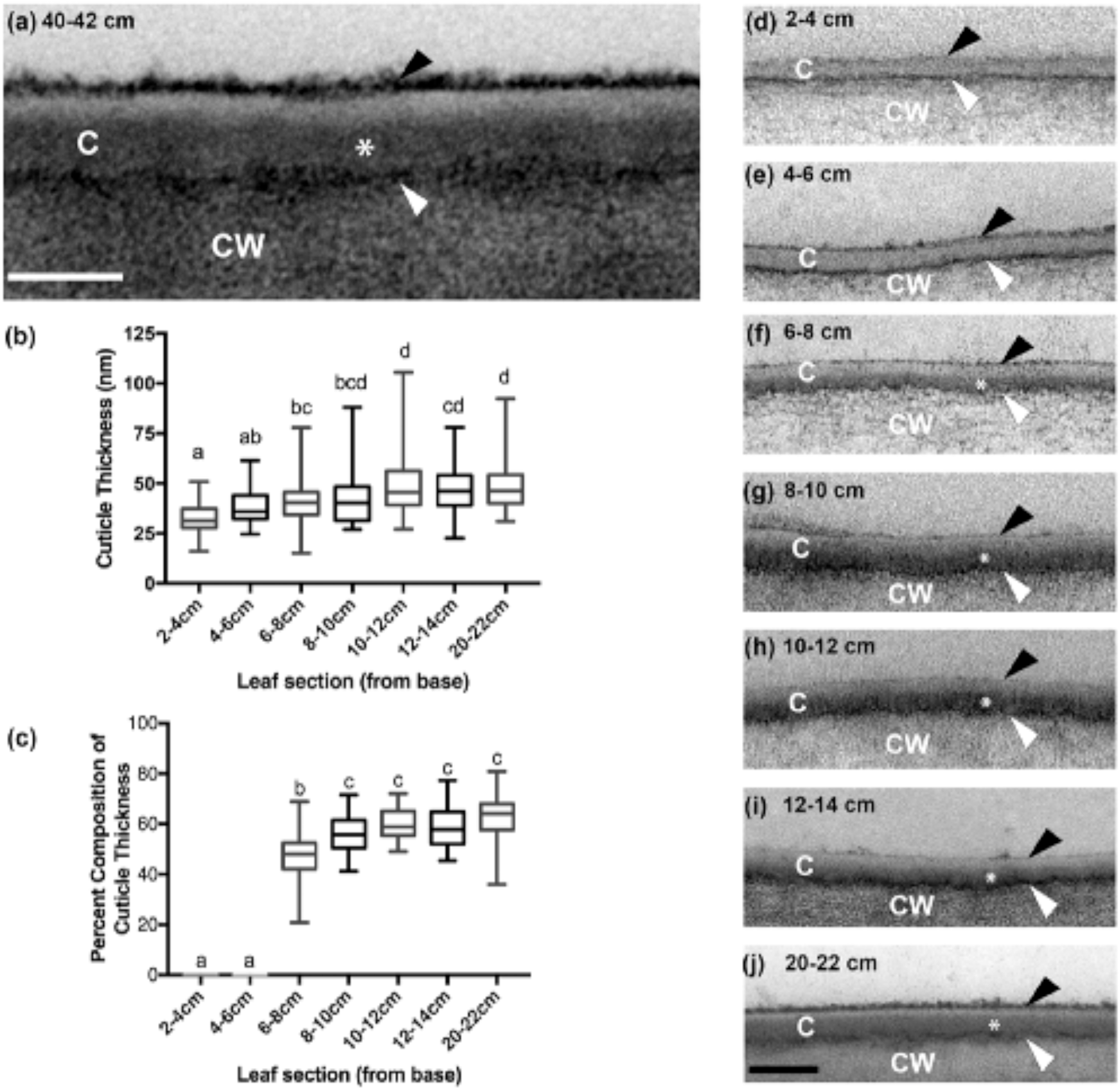
B73 leaf pavement cell cuticle development visualized by transmission electron microscopy. (a) Pavement cell cuticle from a partially expanded leaf 8, 40-42 cm from the base where leaf tissue is mature. Four distinct layers or zones are visible: a thin, darkly-stained layer (white arrowhead) at the interface between the cell wall (CW) and cuticle, dark (white asterisk) and light zones of the cuticle proper (C), and a darkly-stained epicuticular layer (black arrowhead). Scale bar = 40 nm. (b) Thickness of pavement cell cuticles at the indicated positions along the developmental gradient of partially expanded B73 leaf 8. (c) Percent of cuticle thickness at indicated positions occupied by the dark-staining inner layer of the cuticle proper. In (b) and (c), lower case letters indicate significance groups identified by one-way ANOVA with Tukey multiple comparisons post-test. (d-j) Representative images of pavement cell cuticles at the indicated positions along the developmental gradient of partially expanded B73 leaf 8. Scale bar = 100 nm.

**Figure 6.**
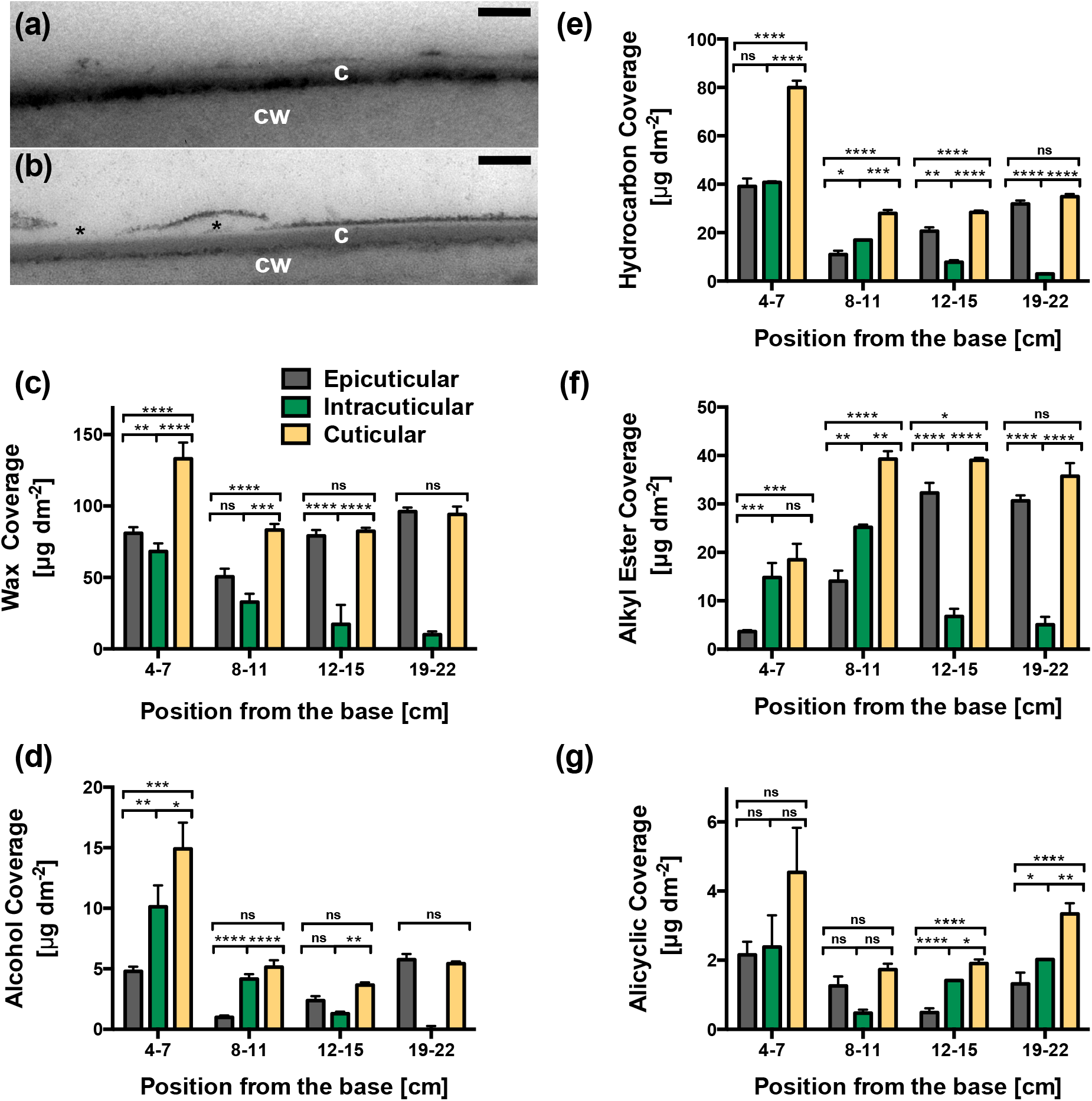
(a) Mature adult maize leaf stripped with gum arabic: most of the dark-staining external layer disappears. (b) Mature adult maize leaves not stripped with gum arabic: the dark-staining external layer of the cuticle becomes detached at places (asterisks), demonstrating a loose association of this layer with the rest of the cuticle. C, cuticle. CW, cell wall. Bars in (a) and (b) = 100 nm. (c-g) Epicuticular, intracuticular and total wax composition in four segments — between 4 and 22 cm from the base— of the maize leaf 8 developmental gradient. Epicuticular waxes removed by adhesion to, and subsequent chloroform extraction from, gum arabic were compared to total wax extracted with chloroform from a matched sample; intracuticular waxes were inferred by calculating the difference between epicuticular vs. total waxes. Total wax coverage (c) and more abundant wax classes, namely alcohols (d), hydrocarbons (e), alkyl esters (f), and alicyclics (g) are shown for each fraction. Means of 3 replicates for chloroform-extracted waxes and of 6 replicates (from 3 adaxial and 3 abaxial) for epicuticular waxes and SE are reported. Statistical analysis was performed with GraphPad Prism, using a one-way ANOVA with Tukey’s multiple comparison post-test. Results were designated significant when the *P*-value (*P*) *<* 0.05 (*, *P <* 0.05; **, *P <* 0.01; ***, *P <* 0.001; ****, *P <* 0.0001; ns, not significant).

We next examined the developmental origins of mature cuticle ultrastructure, by TEM-imaging of pavement cell cuticles at a series of 2-cm intervals along the developmental gradient of a partially expanded adult maize leaf 8. After finding no ultrastructural differences between adaxial and abaxial pavement cell cuticles in the 20-22 cm interval (Fig. S7), we did not attempt to distinguish adaxial and abaxial cuticles in earlier intervals. Cuticle thickness increased by 33% across our developmental gradient, from approximately 30 to 40 nm and reaching this final thickness ~10 cm from the leaf base (Fig. 5b). At the youngest intervals examined (2-4 cm and 4-6 cm from the leaf base), both the wall/cuticle interface and the epicuticular layer were already present, however the cuticle proper was not yet differentiated into darker and lighter zones (Fig. 5d and 5e). The darker zone of the cuticle proper first appeared 6-8 cm from the leaf base (white asterisk, Fig. 5f), reaching its maximum thickness (about two thirds of the thickness of the entire cuticle) approximately 10 cm from the leaf base (Fig. 5g-j and 5c). Prior studies of cuticle ontogeny have consistently shown that dark- and light-staining zones within the cuticle proper emerge during cell expansion (which in our system is completed within 10-12 cm of the leaf base), whereas the cuticular layer emerges later, after completion of cell expansion, in concert with substantial deposition of cutin (e.g., Riederer and Schönherr, 1988; reviewed, Jeffree, 2006). We observed no significant changes in the thickness or appearance of any cuticle zones or layers, or of the cuticle as a whole, beyond the 10-12 cm interval (Fig. 3b,c,g-j), supporting the conclusion that adult maize leaf pavement cells have no cuticular layer. Thus, the ultrastructural features of the mature pavement cell cuticle are established at the same time the cuticle as a whole reaches functional maturity as a barrier to hydrophilic molecules and cell expansion is completed,

### Most cuticular wax of the mature adult maize leaf is epicuticular

To compare the composition of epicuticular and intracuticular waxes and how they change during cuticle ontogeny, epicuticular waxes were mechanically removed with gum arabic from both adaxial and abaxial surfaces (Jetter & Schaffer, 2001) and extracted from gum arabic with chloroform. Epicuticular waxes recovered in this way were compared to total chloroform-extractable cuticular wax; the difference was considered to represent intracuticular wax. For this experiment, 3-cm-long segments of the leaf 8 gradient were analyzed at 4-7, 8-11, 12-15 and 19-22 cm from the base (Fig. 6 c-g). This analysis showed a similar abundance of total waxes (and hydrocarbons, the most abundant wax type at this stage) in both layers between 4 and 11 cm (Fig. 6c and 6e), with a subsequent increase in the relative abundance of epicuticular waxes. By 19-22 cm from the leaf base, the vast majority of extractable waxes were found in the epicuticular fraction (Fig. 6c-f). Alicyclic waxes were found in both epicuticular and intracuticular fractions at all stages analyzed, with similar abundance in both fractions at maturity (19-22cm from the leaf base; Fig. 6g). Consideration of these results in relation to those presented earlier suggests that the bulk of the ultrastructurally-defined cuticle proper consists of wax (predominantly hydrocarbons) at the earliest developmental stages analyzed, but at maturity it is composed mostly of cutin. Thus, functional maturation of the cuticle as a water barrier is associated with a decrease in the amount of intracuticular wax as well as a compositional shift in its wax content concomitant with an increase in cutin.

## Discussion

The cuticle protects shoot tissues from water loss under conditions where stomata are closed, and are thus be a promising target for improving drought tolerance. Cuticle modification through breeding or transgenic strategies is hampered by lack of knowledge of what compositional or structural characteristics of the cuticle are most important for its water barrier properties, and to what extent the determinants are species- or tissue-specific. In this study, we investigated cuticle features related to water permeability by comparing cuticle composition, ultrastructure, and water barrier properties at a series of stages of maize leaf development. Our study is one of the first to characterize cuticles of adult maize leaves, the leaf type determining most of the agronomically significant leaf-related traits in maize.

The development of monocot leaves in a tip-to-base gradient where the youngest tissues are tightly wrapped inside a whorl of older leaves protecting them from desiccation, while older tissues are exposed to the air, provides a unique opportunity to query the relationship between cuticle biogenesis and its acquisition of water barrier properties. Consistent with prior studies focused on juvenile leaves of barley (Richardson *et al.*, 2007) and maize (Hachez *et al.*, 2008), we found that the water barrier property of developing adult maize leaves is established around the same time cell expansion is completed (~10 cm from the leaf base), well before leaves emerge from the whorl (the “point of emergence”, ~17 cm from the leaf base). However, in contrast to findings for developing leaves of juvenile barley plants (Richardson *et al.*, 2007) and leeks (Rhee *et al.*, 1998) where little or no cuticular wax was detected prior to cessation of cell expansion, we found that waxes are already abundant at the earliest stages analyzed in developing maize leaves, where they might have a role in maintaining organ separation, although this function is generally attributed to cutin (reviewed by Fich *et al*., 2016). Our results also contrast with those reported for inbred B73 silk waxes, where 3-fold more hydrocarbons are present on emerged silks compared to encased silk portions (Perera *et al.*, 2010). Thus, there seems to be variation between species, leaf and/or tissue types with respect to the timing of wax deposition during development of monocot organs developing in a tip-to-base gradient.

In spite of the near-constant overall abundance of waxes, wax composition varies dramatically across the adult maize leaf developmental gradient. At 2-4 cm from the leaf base, waxes were composed almost entirely of alkanes and alkenes, which gradually decreased to ~20% of total wax by the time the water barrier is established at ~10 cm. Concomitantly, wax esters increased from nearly undetectable levels initially to become the predominant wax class (~60% of the total) at ~10 cm. Fatty alcohols, free fatty acids and aldehydes remain minor components throughout the developmental period analyzed.

Our results for mature adult maize leaf wax composition are in broad agreement with an earlier study (Bianchi *et al.*, 1984) and show considerable differences from juvenile leaves of maize (e.g., Bianchi *et al.*, 1978) and barley (Richardson *et al.*, 2005), where waxes are dominated by fatty alcohols, with low proportions of aldehydes, alkanes, and esters (Bianchi *et al.*, 1978). Moreover, since previously published scanning EM images reveal no epicuticular wax crystals on the adult leaf surface (e.g., Sylvester & Smith, 2009), we conclude that the epicuticular wax layer we observed via TEM consists of an amorphous film without crystalline structure. While waxes (particularly fatty alcohols and wax esters) are predominantly intracuticular before and during establishment of the water barrier (4-7 and 8-11 cm from the leaf base), at later stages almost all extractable waxes are epicuticular. Maize appears to be unusual in this regard, since most studies comparing epi- and intracuticular waxes in mature leaves of various dicot species show substantial proportions of wax in the intracuticular fraction (e.g. Jetter & Riederer, 2016; Zeisler-Diehl *et al.*, 2018). Although the composition of epicuticular wax changes considerably over the timecourse of adult maize leaf development, its appearance in TEM does not vary. This is consistent with the expectation that saturated aliphatic molecules, whose relative proportions in epicuticular wax are changing, do not react with osmium tetroxide (Cheng et al., 2009; Shumborski et al., 2016). The osmiophilic character of the epicuticular wax film we observe via TEM, which has been observed on leaf surfaces of a variety of other species such as *Hedera helix* (Viougeas *et al.*, 1995) and *Clivia miniata* (Merida *et al.*, 1981) may reflect the presence of unsaturated alicyclic molecules present at developmental stages analyzed, predominantly sterols.

Within the alkane homolog series, we observed a dramatic shift in the distribution of chain lengths over the course of cuticle ontogeny: the blend dominated by C_21_ and C_23_ alkanes 2-4 cm from the leaf base gradually shifts to one dominated by ≥29 carbon alkanes at the point of leaf emergence from the whorl (16-18 cm), remaining similar thereafter. This shift is achieved via reductions in the amounts of most alkanes between 6-12 cm from the leaf base (likely resulting from synthesis not keeping up with rapid cell expansion occurring in this interval) followed by deposition of C_29_-C_37_ alkanes after completion of cell expansion. Shifts from shorter to longer hydrocarbon chain lengths have been previously reported as a feature of cuticle maturation in dicot leaves (Jenks *et al.*, 1996; Jetter & Schaffer, 2001; Busta *et al.*, 2017). In developing Arabidopsis leaves, this shift is associated with increase in the expression level of CER6, a fatty acid elongase needed to produce >C28 hydrocarbons in vitro and in vivo (Busta *et al.*, 2017, Haslam *et al.*, 2012). The functional significance of the shift toward longer hydrocarbon chain lengths is unclear and not apparently related to establishment of the cuticular water barrier property in *Arabidopsis* leaves, which are already exposed to the air prior to the shift. Similarly in maize, we found that accumulation of >30 carbon alkanes occurred after establishment of the water barrier in adult maize leaf cuticles at ~10 cm from the leaf base.

Among the wax compositional changes we observed across the timecourse of adult maize leaf development, only the timing of wax ester accumulation correlates with establishment of the water barrier property of the adult maize leaf cuticle. Thus, our findings suggest a possible role for wax esters in protecting adult maize leaves from dehydration. Consistent with this conclusion, recent work has demonstrated that although wax esters are a very minor component of cuticular waxes in Arabidopsis, reduction of this wax type in *wsd1* wax ester synthase mutants increases drought sensitivity and cuticular permeability (Patwari *et al.*, 2019). However, it is unclear whether these phenotypic effects reflect a key role for wax esters in cuticular impermeabilty, or are due to changes in stomatal density also observed in *wsd1* mutants. The structure of wax esters consisting of two long hydrocarbon molecules linked by a central ester bond might facilitate a parallel arrangement of hydrocarbon tails that can become organized into crystalline domains within the cuticle that efficiently exclude water (Riederer & Schreiber, 2001; Buschhaus & Jetter, 2012). However, wax ester abundance across the maize leaf developmental timecourse increases only in the epicuticular fraction, while decreasing in the intracuticular fraction. Epicuticular wax has been found to play a minor role, if any, in cuticle impermeabilty to water (Jetter & Riederer, 2016; Zeisler-Diehl *et al.*, 2018). The role of wax esters in resistance to dehydration in adult maize leaves, and the relative contributions of epi-vs. intracuticular wax esters, remains to be investigated.

We found that cutin is present at very low abundance 2-6 cm from the leaf base, gradually increasing in abundance from there to the 16-18 cm interval where leaves emerge from the whorl and are exposed to the air. Unlike the shifts in wax composition observed along the adult maize leaf developmental gradient, we observed a relatively constant ratio between the major cutin monomers at all stages analyzed (approx. 2:1 ratio of ω-hydroxy fatty acids to α,ω-dicarboxylic acids), however it is not possible to determine whether this reflects homogeneity in cutin architecture over time. Our findings contrast with those for developing juvenile barley leaves, where no further cutin deposition was observed beyond the point where cell expansion was completed (Richardson *et al.*, 2007) but are consistent with findings for developing leaves of *Clivia miniata* (a monocot), where cutin deposition continued well beyond the cessation of cell expansion (Riederer & Schönherr, 1988). In contrast to *Clivia miniata*, where an increase in cuticle thickness parallels the accumulation of cutin, we observed no increase in pavement cell cuticle thickness beyond the 8-10 cm stage. Notably, bulliform and stomatal cells in adult maize leaves have much thicker cuticles. Consequently, the increase in cutin we observe from 10-22 cm from the leaf base is likely due to selective accumulation of cutin in the cuticles of those cells or other specialized cells we did not analyze, such as macrohairs.

While prior studies have described cuticle ontogeny in ultrastructural terms and/or in relation to lipid composition (e.g., Riederer & Schönherr, 1988), few have attempted to relate these developmental events to the functional maturation of the cuticle. Among the ultrastructural changes we observed across the timecourse of adult maize leaf development, the emergence of a dark-staining inner zone of the cuticle proper coincides with establishment of the water barrier property, appearing initially 6-8 cm from the leaf base and reaching its final thickness and appearance at 8-10 cm. While the emergence of this layer coincides with wax ester accumulation in the cuticle as a whole, wax esters are not osmiophilic and accumulate largely in the epicuticular fraction. The emergence of the dark-staining inner layer of the cuticle proper more likely reflects the deposition of cutin. While cutin alone is not an effective water barrier, it is thought to provide a scaffold for the deposition of waxes needed to form a functional cuticle. Thus, we hypothesize that although intracuticular waxes are depleted as the cuticle matures, an interaction between cutin and the remaining intracuticular waxes that is established initially between 6 and 10 cm from the leaf base is a key feature of cuticular resistance to water loss in the adult maize leaf.

Developmental analysis of the adult maize leaf cuticle revealed unexpectedly dynamic changes in cuticle composition from early stages where cells are dividing to late stages where leaf tissue is photosynthetically mature and is exposed to air and light. Integration of results from our biochemical, ultrastrutural, and functional analyses suggest important roles for wax esters and an ultrastructurally defined, osmiophilic (likely cutin-rich) layer in protection of leaves from dehydration. These studies provide a foundation for future work focused on analysis of underlying gene expression profiles, as well as comparisons between genotypes, aimed at better understanding of the relationship between cuticle composition and function in adult maize leaves and how functionally significant features of the cuticle are supported by underlying gene expression.

## Acknowledgements

This work was supported by NSF Grant IOS-1444507 and by funding from the Canada Research Chairs program and the Canada Foundation for Innovation (CFI-LOF-31502) to I.M. TEM work was conducted at the Cellular and Molecular Medicine Electron microscopy core facility at UCSD, which is supported in part by National Institutes of Health Award number S10OD023527. We thank Joshua Chan (UCSD) for help devising the TBO quantification method.

## Author Contributions

LGS, IM and MJS designed and supervised the research; RB, SM, MV, PQ, AS, CC and MM conducted the research and created the figures; IM and LGS wrote the paper with input from all authors.

## Supplementary data

Additional Supplementary Data may be found online in the Supplementary Data section at the end of the article.

**Fig. S1** Optimization of wax extraction method for mature maize leaf cuticles.

**Fig. S2** Calibration curves for wax ester standards.

**Fig. S3** Identification of monoalkenes in maize leaf wax mixtures.

**Fig. S4** Changes in the alicyclic compound composition in the developmental gradient of mature maize leaves.

**Fig. S5** Comparison of polyester monomers released by transesterification of isolated epidermis and whole leaf tissues.

**Fig. S6** Comparison of cuticles of different cells types in the adult maize leaf epidermis at maturity.

**Fig. S7** Comparison of adaxial and abaxial surfaces of the adult maize leaf.

**Table S1.** Developing maize leaf 8 cuticular wax composition.

**Table S2.** Relative isomeric composition of saturated esters isolated from maize mature leaf cuticular wax and mass spectral data.

**Table S3** Relative cutin monomer composition along the developing maize leaf 8.

**Methods S1** Alkene double bond position analysis.

**Methods S2** Determination of wax ester calibration response factors.

**Methods S3** Enzymatic Isolation of Abaxial and Adaxial Cuticles.

**Notes S1** References cited in Methods S1-S3 that are not included in the main text.

