## Supplementary Data for "Changes in lipid composition and ultrastructure associated with functional maturation of the cuticle during adult maize leaf development"

The following Supplementary data is available for this article:

**Fig. S1** Optimization of wax extraction method for mature maize leaf cuticles.

**Fig. S2** Calibration curves for wax ester standards.

**Fig. S3** Identification of monoalkenes in maize leaf wax mixtures.

**Fig. S4** Changes in the alicyclic compound composition in the developmental gradient of mature maize leaves.

**Fig. S5** Comparison of polyester monomers released by transesterification of isolated epidermis and whole leaf tissues.

**Fig. S6** Comparison of cuticles of different cells types in the adult maize leaf epidermis at maturity.

**Fig. S7** Comparison of adaxial and abaxial surfaces of the adult maize leaf.

**Table S1.** Developing maize leaf 8 cuticular wax composition.

**Table S2.** Relative isomeric composition of saturated esters isolated from maize mature leaf cuticular wax and mass spectral data.

**Table S3** Relative cutin monomer composition along the developing maize leaf 8.

**Methods S1** Alkene double bond position analysis.

**Methods S2** Determination of wax ester calibration response factors**.**

**Methods S3** Enzymatic Isolation of Abaxial and Adaxial Cuticles.

**Notes S1** References cited in Methods S1-S3 that are not included in the main text..

**Fig. S1 Optimization of wax extraction method for mature maize leaf cuticles.** (a) Recovery of fatty alcohols, free fatty acids, alkanes/alkenes, aldehydes, alkyl esters and total waxes extracted with three solvents of different polarity. (b) Comparison of the amounts of wax classes and total waxes after 0.5, 1, 2 and 5 min of extraction with chloroform. Means of 4 replicates +/- SE values are reported. No statistically significant differences in recovery of total waxes as determined by one-way ANOVA with Tukey's HSD test ($F_{2,9} = 0.703$, $p = 0.520$) or individual wax classes ($F_{2,9} < 3.465$, $p > 0.064$) were found among the three different solvents investigated. Similarly, different extraction time lengths for chloroform yielded comparable amounts of total waxes (Games-Howell, $p = 0.721$) and individual wax component groups (ANOVA $F_{3,12} < 3.374$, $p > 0.55$).

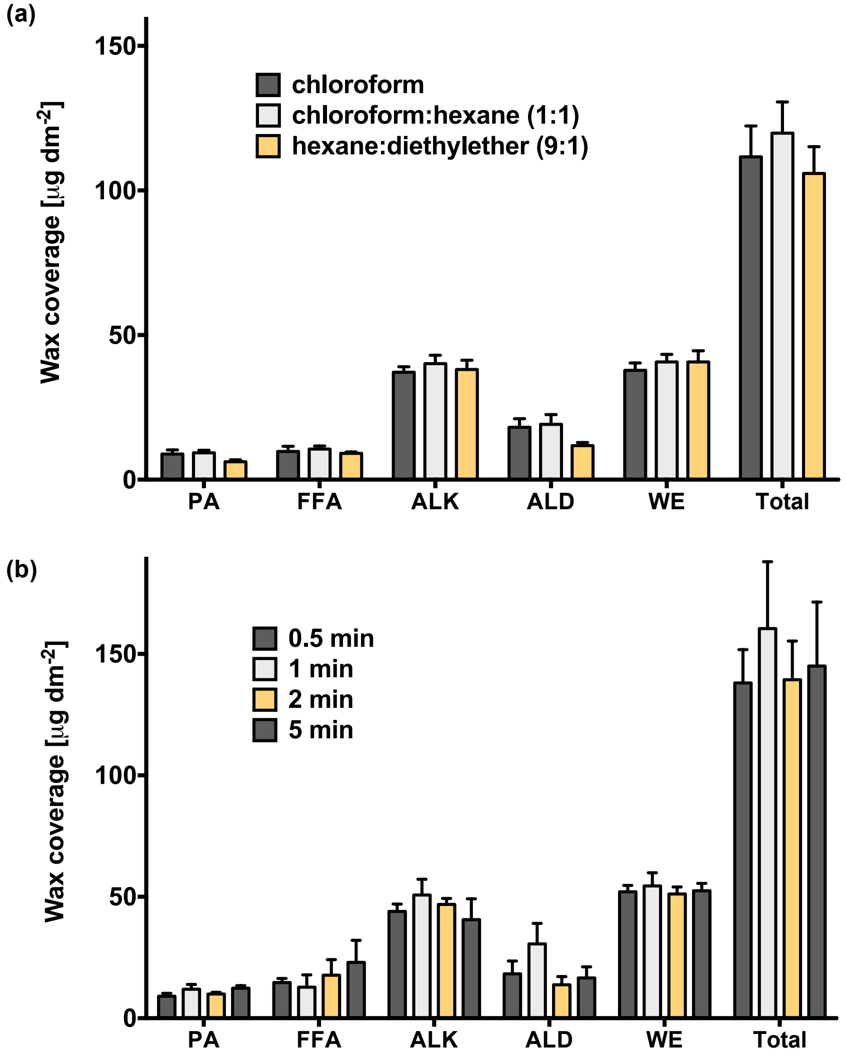

**Fig. S2 . Calibration curves for wax ester standards.** (a-f) Third-order polynomial calibration curves for even-chain saturated wax ester (WE) standards with chain lengths $C_{38}$ - $C_{48}$. Goodness of fit: all curves presented $R^2 > 0.98$ in the ranges of concentration studied, 0 μg - 60 μg (*n* = 3).

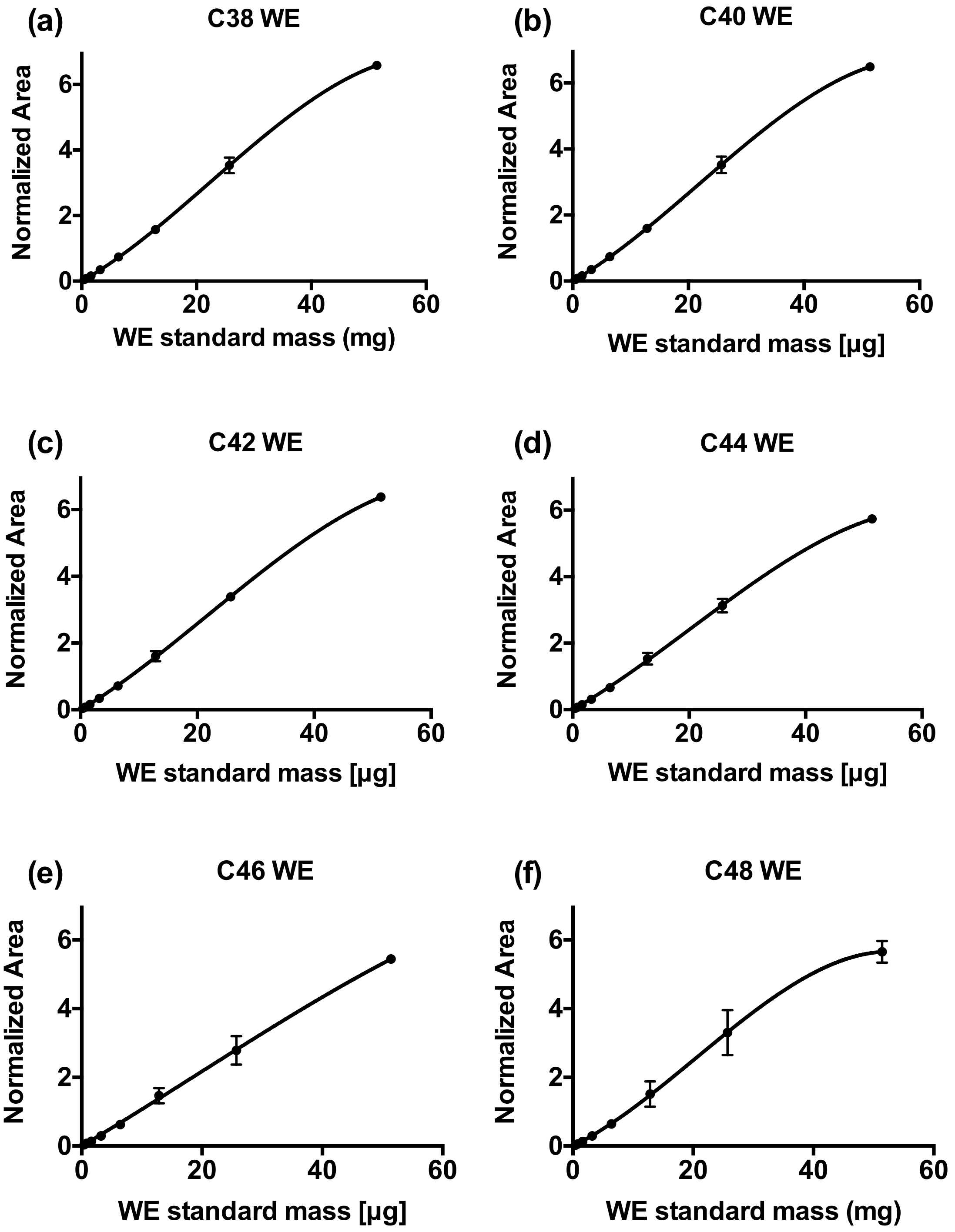

**Fig. S3 Identification of monoalkenes in maize leaf wax mixtures.** Reaction of the alkenes with dimethyl disulfide resulted in adducts that were analyzed by GC-MS.

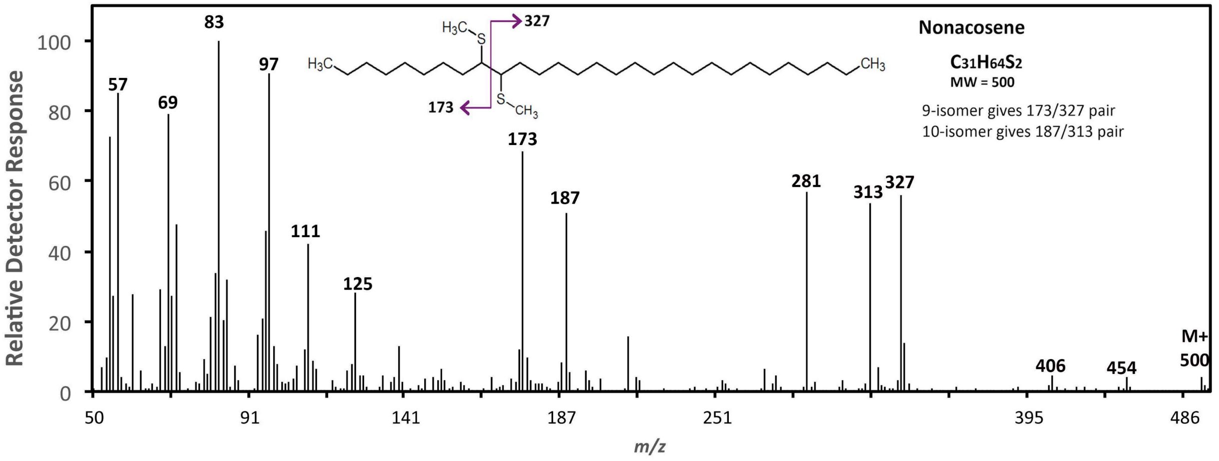


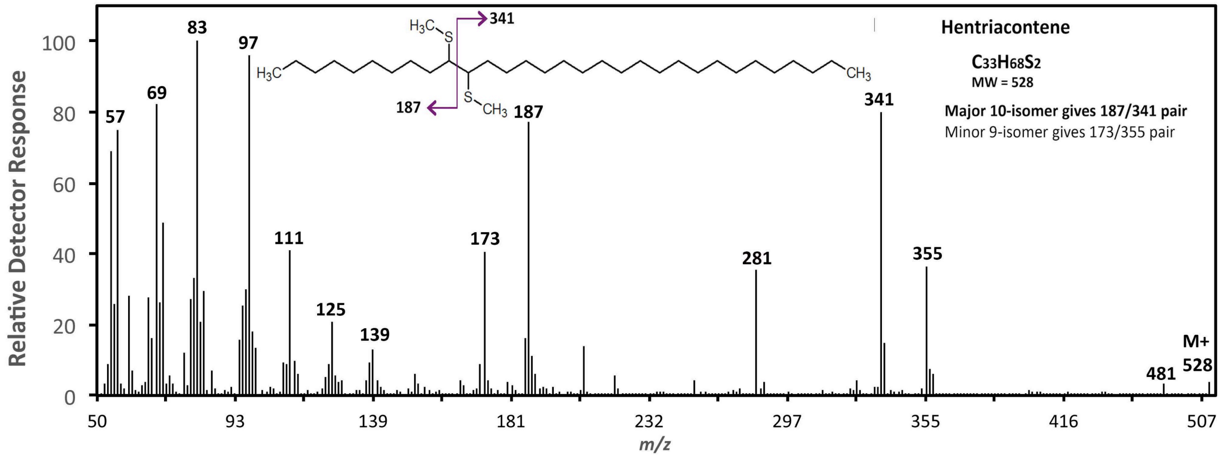

**Fig. S4 Alicyclic wax constituent composition along the developmental gradient of mature maize leaves.** Averages of four independent replicates and SE are reported.

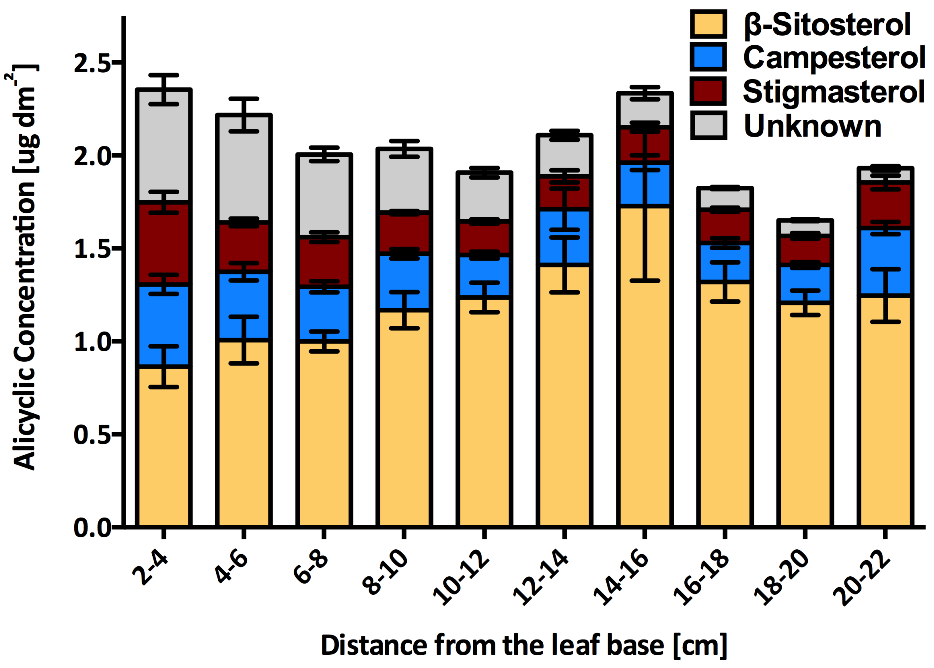

**Fig. S5 Comparison of polyester monomers released by transesterification of isolated epidermis and whole leaf tissues.** Means of 3 replicates +/- SE values are reported. DCAs: dicarboxylic acids; FAs: fatty acids; HCAs: hydroxycinnamic acids; OHFAs: hydroxy fatty acids.

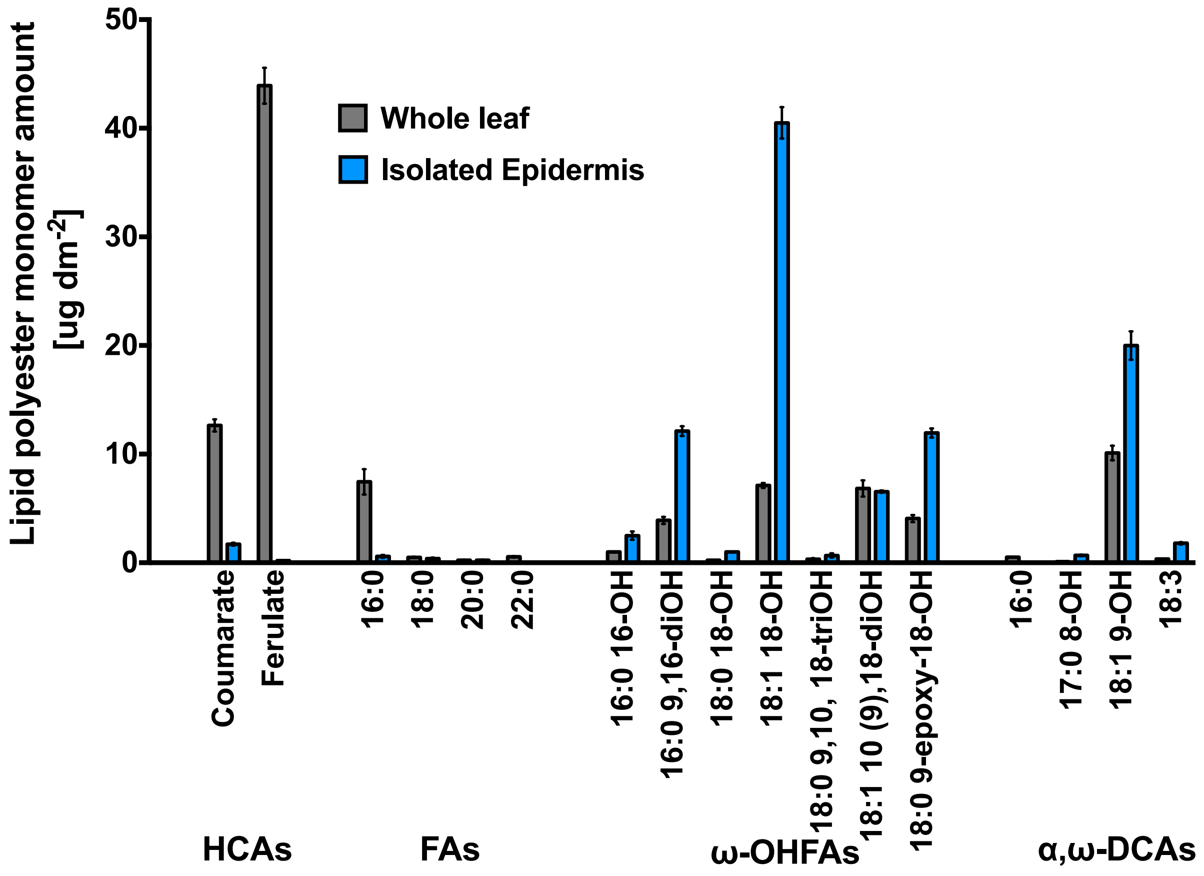

**Fig. S6 Comparison of cuticles of different cells types in the adult maize leaf epidermis at maturity.** (a) pavement cell; (b) bulliform cell; (c) guard cell; (d) stomatal subsidiary cell. White lines demarcate extent of cuticle, which is much thicker in the three specialized cell types shown compared to pavement cells. Bar in C = 200 nm and applies to all images (all are shown at the same magnification).

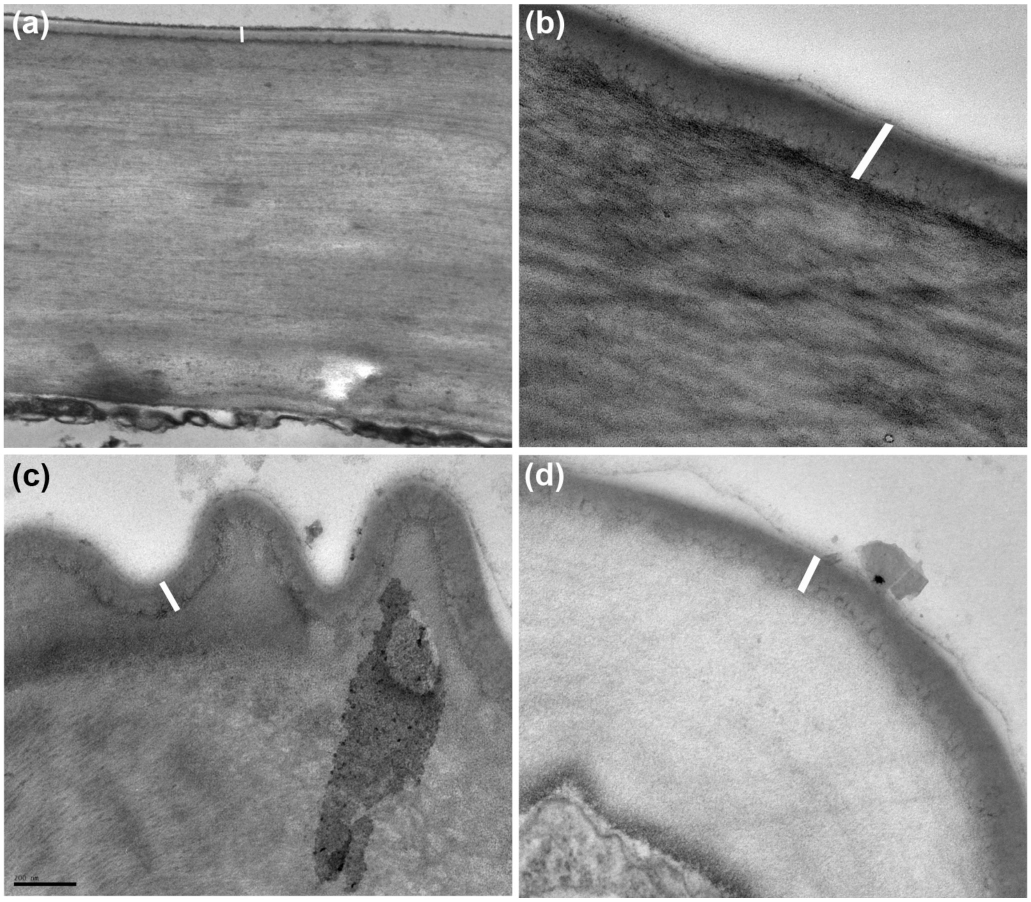

**Fig. S7 Comparison of adaxial and abaxial surfaces of the adult maize leaf.** Adaxial pavement cell cuticle of adult B73 leaf (a) and abaxial pavement cell cuticle of adult B73 leaf (b). Measurements of adaxial and abaxial cuticle thickness from adult B73 pavement cells (c). CW, cell wall; C, cuticle. Scale bar 100nm.

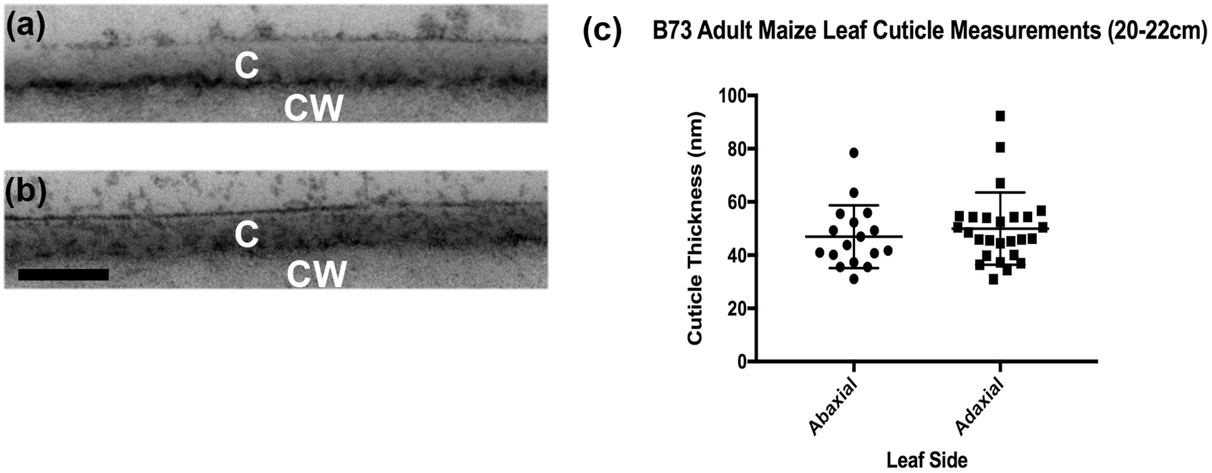


**Table S1** Developing maize leaf 8 cuticular wax composition.

**Table S2** Relative isomeric composition of saturated esters isolated from maize mature leaf cuticular wax and mass spectral data.

**Table S3** Relative cutin monomer composition along the developing maize leaf 8.

**Methods S1** Alkene double bond position analysis.

Wax was extracted by chloroform immersion from the first 6 cm from the base of the leaf, the region in which alkenes predominate. Chloroform was evaporated under $N_2$, and the residual waxes were suspended in hexanes. DMDS adducts of the monounsaturated 29:1 and 31:1 alkenes were formed according to published methods (Dunkelblum et al., 1985; Nichols et al., 1986). Briefly; 100 μL DMDS and 50 μL 6% iodine (w/v) in diethyl ether were added to a 50 μL aliquot of the wax extract in a tightly capped glass tube and heated at 32 °C for 40h. After cooling the reaction mixture, 3.5 mL hexane and 3.5 mL 5% (w/v) aqueous sodium thiosulfate were added and mixed. The organic phase was transferred to a new tube. The aqueous phase was re-extracted with 4:1 hexane/chloroform to increase yield. The combined organic phases were dried under $N_2$ and analyzed by GC-MS as described in Materials and Methods.

**Methods S2**. Determination of wax ester calibration response factors

Calibration curves for wax esters (WEs) with even-chain standards between $C_{38}$ and $C_{48}$ were generated using commercially available standards (Nu-Check Prep Inc., Elysian, MN) following the protocol described by Razeq et al. (2014) with small modifications (samples were spiked a 5 μg of *n*-tetracosane), and using the same GC conditions described in Materials and Methods for wax analysis (Figure S2). These calibration curves were generated using Prism 6 software package (GraphPad Software, Inc., La Jolla, CA, USA). Each wax ester amount was interpolated from their respective calibration curves, with the exception of $C_{50}$ and longer WEs, for which the $C_{48}$ calibration curve was used to estimate their loads in samples.

**Methods S3** Enzymatic Isolation of Abaxial and Adaxial Cuticles

Immediately before use, pectinase 20 U/ml and cellulase 24 U/ml (Marga et al., 2001) were dissolved in 20 mM citrate buffer at pH 3.0 containing 1 mM sodium azide (Schonherr and Riederer, 1986; Franke et al., 2005). The midrib and 2 mm of the outer margin of the leaf were removed from the 14 to 18 cm portion of leaf 8. Each segment was placed in a petri dish, applying a moderate vacuum in a closed chamber for the first 5 min, and then incubated with 15 ml of the hydrolytic enzyme solution at 30 °C for 4-6 d. The separated, floating epidermis/cuticle was observed examined under the microscope to confirm that these did not contain any veins attached. Separated cuticles/epidermis samples and leaf controls not exposed to enzyme treatment were placed in glass, screw-cap tubes containing isopropanol. Each sample was processed for cutin analysis as described in Materials and Methods.

**Notes S1** References cited in Methods S1-S3 that are not included in the main text.

**Dunkelblum E, Tan SH, Silk PJ. 1985.** Double-bond location in monounsaturated fatty acids by dimethyl disulfide derivatization and mass spectrometry: application to analysis of fatty acids in pheromone glands of four Lepidoptera. *Journal of Chemical Ecology* **11:** 265–277.

**Franke R, Briesen I, Wojciechowski T, Faust A, Yephremov A, Nawrath C, Schreiber L. 2005.** Apoplastic polyesters in Arabidopsis surface tissues--a typical suberin and a particular cutin. *Phytochemistry* **66:** 2643-58.

**Marga F, Pesacreta TC, Hasenstein KH. 2001.** Biochemical analysis of elastic and rigid cuticles of *Cirsium horridulum*. *Planta* **213:** 841-8.

**Nichols PD, Guckert JB, White DC.1986.** Determination of monounsaturated fatty acid double-bond position and geometry for microbial monocultures and complex consortia by capillary GC-MS of their dimethyl disulphide adducts. *Journal of Microbiological Methods* **5:** 49-55.

**Schönherr J, Riederer M. 1986.** Plant cuticles sorb lipophilic compounds during enzymatic isolation. *Plant, Cell & Environment* **4:** 349-354.
