## Supplemental Table 1 for "Changes in lipid composition and ultrastructure associated with functional maturation of the cuticle during adult maize leaf development"

**Table S1. Developing maize leaf 8 cuticular wax composition. (Values reported as mass %)**

| Molecular class | Carbon # : double bond # or name | Position along the leaf #8 blade (cm from the base) | | | | | | | | | | | | | | | | | | | |
|---|---|---|---|---|---|---|---|---|---|---|---|---|---|---|---|---|---|---|---|---|---|
| | | 2-4 | | 4-6 | | 6-8 | | 8-10 | | 10-12 | | 12-14 | | 14-16 | | 16-18 | | 18-20 | | 20-22 | |
| | | Mean | SE | Mean | SE | Mean | SE | Mean | SE | Mean | SE | Mean | SE | Mean | SE | Mean | SE | Mean | SE | Mean | SE |
| **Fatty alcohols** | 24:0 | 0.18 | 0.02 | 2.42 | 0.38 | 3.83 | 0.27 | 1.19 | 0.13 | 0.43 | 0.04 | 0.28 | 0.02 | 0.33 | 0.09 | 0.32 | 0.03 | 0.34 | 0.02 | 0.36 | 0.03 |
| | 26:0 | 0.14 | 0.02 | 1.50 | 0.21 | 3.05 | 0.17 | 2.06 | 0.14 | 1.09 | 0.10 | 0.86 | 0.14 | 0.79 | 0.20 | 0.77 | 0.04 | 1.15 | 0.10 | 1.47 | 0.09 |
| | 28:0 | 0.09 | 0.01 | 0.75 | 0.09 | 1.01 | 0.06 | 0.63 | 0.03 | 0.38 | 0.03 | 0.34 | 0.06 | 0.43 | 0.14 | 0.38 | 0.01 | 0.63 | 0.02 | 0.99 | 0.12 |
| | 30:0 | 0.04 | 0.01 | 0.93 | 0.08 | 1.05 | 0.05 | 0.73 | 0.05 | 0.62 | 0.04 | 0.65 | 0.06 | 0.84 | 0.18 | 0.76 | 0.03 | 1.11 | 0.06 | 1.40 | 0.17 |
| | 32:0 | 0.37 | 0.01 | 1.74 | 0.09 | 2.30 | 0.11 | 1.71 | 0.10 | 1.34 | 0.09 | 1.51 | 0.15 | 1.96 | 0.23 | 1.70 | 0.07 | 1.85 | 0.15 | 1.84 | 0.20 |
| | 34:0 | 0.20 | 0.00 | 0.71 | 0.05 | 1.42 | 0.11 | 0.92 | 0.06 | 0.75 | 0.04 | 0.72 | 0.04 | 0.88 | 0.10 | 0.77 | 0.02 | 0.87 | 0.07 | 0.80 | 0.07 |
| | | **1.01** | **0.09** | **8.06** | **0.89** | **12.66** | **0.78** | **7.24** | **0.51** | **4.61** | **0.34** | **4.36** | **0.47** | **5.22** | **0.94** | **4.71** | **0.20** | **5.96** | **0.41** | **6.86** | **0.68** |
| **Fatty acids** | | | | | | | | | | | | | | | | | | | | | |
| | 16:0 | 2.44 | 0.32 | 1.83 | 0.06 | 1.92 | 0.11 | 2.02 | 0.15 | 2.29 | 0.11 | 2.10 | 0.04 | 2.31 | 0.17 | 2.35 | 0.11 | 2.19 | 0.09 | 2.42 | 0.30 |
| | 18:0 | 2.53 | 0.25 | 2.21 | 0.10 | 2.17 | 0.08 | 2.59 | 0.19 | 2.84 | 0.14 | 2.73 | 0.14 | 2.77 | 0.24 | 2.77 | 0.14 | 2.62 | 0.09 | 2.55 | 0.13 |
| | | **4.96** | **0.57** | **4.04** | **0.16** | **4.09** | **0.19** | **4.60** | **0.34** | **5.13** | **0.25** | **4.84** | **0.19** | **5.08** | **0.42** | **5.11** | **0.25** | **4.82** | **0.18** | **4.97** | **0.43** |
| **Hydrocarbons** | | | | | | | | | | | | | | | | | | | | | |
| | 21:0 | 24.49 | 1.18 | 8.82 | 0.19 | 2.62 | 0.15 | 0.71 | 0.09 | 0.26 | 0.02 | 0.17 | 0.08 | 0.08 | 0.01 | 0.04 | 0.01 | 0.02 | 0.01 | 0.02 | 0.00 |
| | 23:0 | 16.78 | 0.63 | 14.96 | 0.45 | 8.93 | 0.35 | 5.17 | 0.21 | 3.30 | 0.08 | 1.97 | 0.22 | 0.67 | 0.02 | 0.20 | 0.01 | 0.09 | 0.01 | 0.06 | 0.00 |
| | 25:0 | 3.20 | 0.10 | 3.38 | 0.10 | 2.36 | 0.08 | 1.73 | 0.04 | 1.41 | 0.02 | 1.49 | 0.43 | 0.78 | 0.03 | 0.53 | 0.02 | 0.35 | 0.02 | 0.23 | 0.01 |
| | 27:0 | 6.31 | 0.21 | 6.01 | 0.23 | 4.05 | 0.21 | 2.91 | 0.04 | 2.45 | 0.06 | 2.41 | 0.44 | 1.89 | 0.08 | 1.82 | 0.05 | 1.71 | 0.04 | 1.59 | 0.01 |
| | 29:1 | 11.17 | 0.49 | 8.08 | 0.49 | 5.17 | 0.27 | 3.77 | 0.10 | 3.05 | 0.06 | 2.05 | 0.06 | 1.23 | 0.04 | 0.76 | 0.04 | 0.43 | 0.03 | 0.25 | 0.03 |
| | 29:0 | 10.42 | 0.36 | 10.00 | 0.28 | 7.42 | 0.27 | 5.58 | 0.09 | 4.88 | 0.10 | 4.61 | 0.27 | 4.53 | 0.20 | 5.01 | 0.13 | 6.04 | 0.12 | 6.82 | 0.11 |
| | 31:1 | 12.45 | 0.57 | 9.04 | 0.42 | 6.01 | 0.23 | 4.24 | 0.05 | 3.51 | 0.07 | 2.66 | 0.04 | 1.89 | 0.07 | 1.34 | 0.07 | 0.87 | 0.04 | 0.58 | 0.04 |
| | 31:0 | 2.57 | 0.10 | 1.82 | 0.05 | 1.54 | 0.05 | 1.29 | 0.03 | 2.51 | 0.18 | 6.42 | 0.30 | 8.98 | 0.57 | 10.07 | 0.29 | 11.01 | 0.27 | 11.89 | 0.21 |
| | 33:1 | 0.80 | 0.03 | 0.67 | 0.03 | 0.52 | 0.03 | 0.40 | 0.01 | 0.32 | 0.01 | 0.28 | 0.01 | 0.27 | 0.01 | 0.23 | 0.02 | 0.21 | 0.00 | 0.19 | 0.01 |
| | 33:0 | 0.12 | 0.01 | 0.07 | 0.00 | 0.06 | 0.01 | 0.20 | 0.03 | 1.26 | 0.08 | 2.76 | 0.04 | 3.45 | 0.03 | 3.83 | 0.15 | 4.51 | 0.18 | 5.06 | 0.06 |
| | 35:0 | 0.01 | 0.00 | 0.01 | 0.00 | 0.00 | 0.00 | 0.06 | 0.01 | 0.45 | 0.02 | 0.91 | 0.04 | 1.19 | 0.03 | 1.37 | 0.06 | 1.68 | 0.06 | 1.85 | 0.03 |
| | 37:0 | 0.02 | 0.00 | 0.07 | 0.01 | 0.06 | 0.02 | 0.04 | 0.00 | 0.08 | 0.00 | 0.14 | 0.01 | 0.19 | 0.01 | 0.24 | 0.01 | 0.29 | 0.01 | 0.30 | 0.01 |
| | | **88.34** | **3.69** | **62.93** | **2.26** | **38.75** | **1.65** | **26.10** | **0.70** | **23.49** | **0.70** | **25.86** | **1.94** | **25.15** | **1.11** | **25.45** | **0.86** | **27.21** | **0.79** | **28.84** | **0.54** |
| **Aldehydes** | | | | | | | | | | | | | | | | | | | | | |
| | 26:0 | 0.04 | 0.01 | 0.13 | 0.03 | 0.05 | 0.00 | 0.09 | 0.01 | 0.16 | 0.02 | 0.10 | 0.01 | 0.06 | 0.00 | 0.08 | 0.01 | 0.41 | 0.06 | 0.67 | 0.03 |
| | 28:0 | 0.04 | 0.02 | 0.05 | 0.00 | 0.03 | 0.00 | 0.02 | 0.00 | 0.01 | 0.00 | 0.05 | 0.05 | 0.02 | 0.00 | 0.09 | 0.02 | 0.62 | 0.07 | 1.59 | 0.15 |
| | 30:0 | 0.02 | 0.00 | 0.03 | 0.00 | 0.04 | 0.00 | 0.03 | 0.00 | 0.04 | 0.00 | 0.03 | 0.00 | 0.04 | 0.01 | 0.13 | 0.02 | 0.93 | 0.15 | 2.48 | 0.37 |
| | 32:0 | 0.01 | 0.00 | 0.01 | 0.00 | 0.04 | 0.01 | 0.04 | 0.01 | 0.04 | 0.01 | 0.05 | 0.01 | 0.06 | 0.01 | 0.10 | 0.01 | 0.41 | 0.10 | 0.92 | 0.16 |
| | 34:0 | 0.00 | 0.00 | 0.00 | 0.00 | 0.01 | 0.00 | 0.01 | 0.00 | 0.01 | 0.00 | 0.02 | 0.01 | 0.01 | 0.00 | 0.02 | 0.00 | 0.10 | 0.03 | 0.18 | 0.03 |
| | | **0.11** | **0.03** | **0.23** | **0.04** | **0.17** | **0.01** | **0.20** | **0.02** | **0.26** | **0.02** | **0.26** | **0.08** | **0.19** | **0.02** | **0.42** | **0.05** | **2.47** | **0.40** | **5.85** | **0.74** |
| **Alkyl Esters** | | | | | | | | | | | | | | | | | | | | | |
| | 38:0 | 0.01 | 0.00 | 0.12 | 0.01 | 0.21 | 0.01 | 0.21 | 0.01 | 0.17 | 0.05 | 0.12 | 0.01 | 0.09 | 0.01 | 0.08 | 0.00 | 0.06 | 0.00 | 0.03 | 0.00 |
| | 40:0 | 0.03 | 0.00 | 0.50 | 0.06 | 1.57 | 0.11 | 2.10 | 0.05 | 2.11 | 0.12 | 1.89 | 0.05 | 1.73 | 0.07 | 1.49 | 0.04 | 1.14 | 0.03 | 0.90 | 0.06 |
| | 41:0 | 0.01 | 0.00 | 0.12 | 0.01 | 0.31 | 0.02 | 0.44 | 0.01 | 0.43 | 0.04 | 0.39 | 0.02 | 0.40 | 0.02 | 0.42 | 0.01 | 0.37 | 0.01 | 0.31 | 0.03 |
| | 42:0 | 0.06 | 0.01 | 0.74 | 0.09 | 3.69 | 0.25 | 6.40 | 0.26 | 7.03 | 0.04 | 6.85 | 0.07 | 6.90 | 0.21 | 6.77 | 0.19 | 6.14 | 0.08 | 5.62 | 0.18 |
| | 43:0 | 0.01 | 0.00 | 0.09 | 0.01 | 0.26 | 0.02 | 0.43 | 0.01 | 0.49 | 0.11 | 0.50 | 0.01 | 0.53 | 0.03 | 0.54 | 0.02 | 0.51 | 0.02 | 0.46 | 0.04 |
| | 44:0 | 0.28 | 0.01 | 1.61 | 0.17 | 7.15 | 0.50 | 15.37 | 0.96 | 18.18 | 0.05 | 17.94 | 0.25 | 17.58 | 0.34 | 17.14 | 0.59 | 15.63 | 0.47 | 14.43 | 0.53 |
| | 45:0 | 0.01 | 0.00 | 0.12 | 0.00 | 0.27 | 0.03 | 0.49 | 0.04 | 0.64 | 0.00 | 0.65 | 0.01 | 0.69 | 0.04 | 0.70 | 0.03 | 0.66 | 0.03 | 0.61 | 0.05 |
| | 46:0 | 0.20 | 0.01 | 1.76 | 0.16 | 4.29 | 0.26 | 7.89 | 0.57 | 9.99 | 0.13 | 10.27 | 0.30 | 10.27 | 0.36 | 10.54 | 0.36 | 9.83 | 0.26 | 8.88 | 0.31 |
| | 47:0 | 0.01 | 0.00 | 0.08 | 0.02 | 0.12 | 0.00 | 0.17 | 0.01 | 0.20 | 0.17 | 0.24 | 0.03 | 0.22 | 0.01 | 0.24 | 0.01 | 0.24 | 0.01 | 0.22 | 0.01 |
| | 48:0 | 0.15 | 0.02 | 1.22 | 0.09 | 1.99 | 0.15 | 2.59 | 0.20 | 2.88 | 0.00 | 2.91 | 0.15 | 2.94 | 0.11 | 3.23 | 0.09 | 3.20 | 0.11 | 2.90 | 0.09 |
| | 49:0 | 0.01 | 0.00 | 1.73 | 0.11 | 2.32 | 0.18 | 2.03 | 0.07 | 1.85 | 0.02 | 1.87 | 0.05 | 1.90 | 0.08 | 2.07 | 0.05 | 2.07 | 0.04 | 1.82 | 0.06 |
| | 50:0 | 0.21 | 0.01 | 0.72 | 0.05 | 0.99 | 0.08 | 0.89 | 0.03 | 0.81 | 0.09 | 0.81 | 0.03 | 0.83 | 0.03 | 0.92 | 0.02 | 0.92 | 0.02 | 0.81 | 0.03 |
| | 52:0 | 0.23 | 0.00 | 2.26 | 0.15 | 4.18 | 0.25 | 4.18 | 0.18 | 3.71 | 0.02 | 3.59 | 0.03 | 3.73 | 0.20 | 3.83 | 0.13 | 3.55 | 0.02 | 3.13 | 0.17 |
| | 54:0 | 0.59 | 0.05 | 4.46 | 0.31 | 7.27 | 0.35 | 8.24 | 0.35 | 7.58 | 0.07 | 6.96 | 0.09 | 6.95 | 0.42 | 7.00 | 0.23 | 6.40 | 0.06 | 5.53 | 0.32 |
| | 56:0 | 0.93 | 0.07 | 5.37 | 0.28 | 6.12 | 0.31 | 6.61 | 0.41 | 6.41 | 0.07 | 5.79 | 0.25 | 5.61 | 0.41 | 5.83 | 0.32 | 5.55 | 0.12 | 4.67 | 0.16 |
| | 58:0 | 0.68 | 0.03 | 1.90 | 0.11 | 1.82 | 0.12 | 1.77 | 0.13 | 1.76 | 0.12 | 1.55 | 0.10 | 1.49 | 0.14 | 1.60 | 0.11 | 1.52 | 0.07 | 1.29 | 0.07 |
| | | **3.43** | **0.21** | **22.80** | **1.64** | **42.55** | **2.64** | **59.81** | **3.30** | **64.23** | **1.09** | **62.34** | **1.45** | **61.87** | **2.46** | **62.39** | **2.20** | **57.79** | **1.37** | **51.64** | **2.11** |
| **Sterols** | | | | | | | | | | | | | | | | | | | | | |
| | β-Sitosterol | 0.79 | 0.10 | 0.89 | 0.11 | 0.89 | 0.05 | 1.18 | 0.10 | 1.47 | 0.09 | 1.57 | 0.16 | 1.84 | 0.43 | 1.39 | 0.11 | 1.28 | 0.07 | 1.19 | 0.14 |
| | Campesterol | 0.40 | 0.05 | 0.32 | 0.04 | 0.26 | 0.03 | 0.31 | 0.03 | 0.27 | 0.02 | 0.33 | 0.12 | 0.25 | 0.04 | 0.22 | 0.03 | 0.22 | 0.02 | 0.35 | 0.03 |
| | Stigmasterol | 0.40 | 0.05 | 0.23 | 0.02 | 0.24 | 0.02 | 0.22 | 0.01 | 0.21 | 0.01 | 0.20 | 0.04 | 0.20 | 0.03 | 0.19 | 0.01 | 0.17 | 0.02 | 0.23 | 0.04 |
| | Non identified | 0.55 | 0.07 | 0.51 | 0.08 | 0.40 | 0.03 | 0.35 | 0.04 | 0.31 | 0.03 | 0.24 | 0.03 | 0.20 | 0.03 | 0.12 | 0.01 | 0.09 | 0.01 | 0.07 | 0.01 |
| | | **2.15** | **0.27** | **1.95** | **0.25** | **1.79** | **0.13** | **2.05** | **0.18** | **2.27** | **0.16** | **2.34** | **0.35** | **2.49** | **0.53** | **1.92** | **0.16** | **1.75** | **0.11** | **1.85** | **0.21** |
| | Total | 100.00 | 4.86 | 100.00 | 5.23 | 100.00 | 5.41 | 100.00 | 5.05 | 100.00 | 2.56 | 100.00 | 4.47 | 100.00 | 5.48 | 100.00 | 3.73 | 100.00 | 3.26 | 100.00 | 4.72 |
