## Supplemental Table 2 for "Changes in lipid composition and ultrastructure associated with functional maturation of the cuticle during adult maize leaf development"

**Table 2.** Relative isomeric composition of saturated esters isolated from maize mature leaf cuticular wax and mass spectral data.

| Wax ester (CN:DB)[a] | Components (acid–alcohol) | % of isomer[b] | SD | $[M]^+$ | $[RCO2H2]^+$ | $[R' – 1]^+$ |
|---|---|---|---|---|---|---|
| **WE 42:0** | 16:0–26:0 | 16.4 | 2.0 | 621 | 257 | 364 |
| | 18:0–24:0 | **33.7** | **2.6** | | 285 | 336 |
| | 20:0–22:0 | **30.7** | **2.6** | | 313 | 308 |
| | 21:0–21:0 | 4.1 | 0.9 | | 327 | 294 |
| | 22:0–20:0 | 7.7 | 3.4 | | 341 | 280 |
| | 23:0–19:0 | 4.5 | 1.4 | | 355 | 266 |
| | *Minor isomers | 2.9 | 1.1 | | | |
| **WE 44:0** | 16:0–28:0 | 2.1 | 0.9 | 649 | 257 | 392 |
| | 18:0–26:0 | **11.5** | **2.4** | | 285 | 364 |
| | 20:0–24:0 | **67.0** | **4.3** | | 313 | 336 |
| | 21:0–23:0 | 1.3 | 0.4 | | 327 | 322 |
| | 22:0–22:0 | **11.1** | **3.7** | | 341 | 308 |
| | 23:0–21:0 | 3.6 | 1.2 | | 355 | 294 |
| | *Minor isomers | 3.4 | 0.9 | | | |
| **WE 46:0** | 16:0–30:0 | 1.8 | 0.1 | 677 | 257 | 420 |
| | 18:0–28:0 | 2.3 | 0.1 | | 285 | 392 |
| | 20:0–26:0 | **62.0** | **2.4** | | 313 | 364 |
| | 21:0–25:0 | 1.8 | 1.5 | | 327 | 350 |
| | 22:0–24:0 | **26.4** | **3.7** | | 341 | 336 |
| | 23:0–23:0 | 2.0 | 0.8 | | 355 | 322 |
| | 24:0–22:0 | 1.0 | 0.3 | | 369 | 308 |
| | *Minor isomers | 2.80 | 0.6 | | | |
| **WE 48:0** | 16:0–32:0 | 3.8 | 0.5 | 705 | 257 | 448 |
| | 18:0–30:0 | 7.0 | 1.2 | | 285 | 420 |
| | 20:0–28:0 | **23.2** | **1.0** | | 313 | 392 |
| | 22:0-26:0 | **51.2** | **1.9** | | 341 | 364 |
| | 24:0-24:0 | **10.6** | **0.6** | | 369 | 336 |
| | *Minor isomers | 4.30 | 1.0 | | | |
| **WE 50:0** | 16:0–34:0 | 6.3 | 2.4 | 733 | 257 | 476 |
| | 18:0–32:0 | **17.3** | **8.6** | | 285 | 448 |
| | 20:0–30:0 | **43.4** | **4.1** | | 313 | 420 |
| | 22:0-28:0 | **12.7** | **20.0** | | 341 | 392 |
| | 24:0-26:0 | **19.5** | **8.4** | | 369 | 364 |
| | *Minor isomers | 0.9 | 0.4 | | | |
| **WE 52:0** | 16:0–36:0 | 5.6 | 1.4 | 761 | 257 | 504 |
| | 18:0–34:0 | 9.3 | 0.0 | | 285 | 476 |
| | 20:0–32:0 | **43.5** | **7.4** | | 313 | 448 |
| | 22:0-30:0 | **36.3** | **6.8** | | 341 | 420 |
| | 26:0-26:0 | 1.3 | 0.0 | | 397 | 364 |
| | *Minor isomers | 4.0 | 1.7 | | | |
| **WE 54:0** | 16:0–38:0 | 2.2 | 0.7 | 789 | 257 | 532 |
| | 18:0–36:0 | 7.3 | 0.9 | | 285 | 504 |
| | 20:0–34:0 | **41.7** | **1.2** | | 313 | 476 |
| | 22:0-32:0 | **45.1** | **1.3** | | 341 | 448 |
| | 30:0-24:0 | 1.2 | 0.2 | | 453 | 336 |
| | *Minor isomers | 2.4 | 1.5 | | | |
| **WE 56:0** | 16:0–40:0 | 1.2 | 1.1 | 817 | 257 | 560 |
| | 18:0–38:0 | 4.0 | 0.9 | | 285 | 532 |
| | 20:0–36:0 | **24.9** | **9.8** | | 313 | 504 |
| | 22:0-34:0 | **50.0** | **1.3** | | 341 | 476 |
| | 24:0-32:0 | **16.2** | **13.0** | | 369 | 448 |
| | 28:0-28:0 | 1.0 | 0.6 | | 425 | 392 |

[a] CN, carbon number of wax ester; DB, number of double bonds; WE, wax ester.

[b] The percentage of a single isomer of a saturated wax ester was calculated based on intensities of the $[RCO2H2]^+$ (acyl moiety) ion from 4 biological replicates.

*Wax ester homologues constituting less than 1% of the mixture were pooled and included in the "minor isomers" category.
