## Supplemental Table 3 for "Changes in lipid composition and ultrastructure associated with functional maturation of the cuticle during adult maize leaf development"

**Table S3.** Relative cutin monomer composition along the developing maize leaf (values reported in Mole %).

| Molecular clas | Carbon # : double bond # | Position along the leaf #8 blade (cm from the base) | | | | | | | | | | | | | | | | | | | |
|---|---|---|---|---|---|---|---|---|---|---|---|---|---|---|---|---|---|---|---|---|---|
| | | 2-4 | | 4-6 | | 6-8 | | 8-10 | | 10-12 | | 12-14 | | 14-16 | | 16-18 | | 18-20 | | 20-22 | |
| | | Mean | SE | Mean | SE | Mean | SE | Mean | SE | Mean | SE | Mean | SE | Mean | SE | Mean | SE | Mean | SE | Mean | SE |
| **Fatty acids** | | | | | | | | | | | | | | | | | | | | | |
| | 18:0 | 28.41 | 2.10 | 21.12 | 5.82 | 9.60 | 2.10 | 4.64 | 0.81 | 2.33 | 0.15 | 2.09 | 0.28 | 1.47 | 0.04 | 1.25 | 0.22 | 1.35 | 0.16 | 1.47 | 0.24 |
| | 20:0 | 9.88 | 2.09 | 11.40 | 1.21 | 5.97 | 0.37 | 2.20 | 0.25 | 1.12 | 0.26 | 1.02 | 0.08 | 0.81 | 0.07 | 0.57 | 0.08 | 0.65 | 0.09 | 0.69 | 0.07 |
| | 22:0 | 7.93 | 0.85 | 9.13 | 1.72 | 6.36 | 0.71 | 2.98 | 0.31 | 1.65 | 0.09 | 1.46 | 0.13 | 1.22 | 0.04 | 1.28 | 0.09 | 1.52 | 0.15 | 1.66 | 0.07 |
| | 24:0 | 0.00 | 0.00 | 0.00 | 0.00 | 0.00 | 0.00 | 0.42 | 0.05 | 0.34 | 0.09 | 0.31 | 0.01 | 0.26 | 0.01 | 0.30 | 0.06 | 0.38 | 0.05 | 0.52 | 0.05 |
| **ω-Hydroxyacids** | | 0.00 | 0.00 | 0.00 | 0.00 | 0.00 | 0.00 | 0.00 | 0.00 | 0.00 | 0.00 | 0.00 | 0.00 | 0.00 | 0.00 | 0.00 | 0.00 | 0.00 | 0.00 | 0.00 | 0.00 |
| | 16:0 16-hydroxy | 0.00 | 0.00 | 0.00 | 0.00 | 5.70 | 0.37 | 5.24 | 0.30 | 4.62 | 0.11 | 4.60 | 0.18 | 3.72 | 0.16 | 2.90 | 0.07 | 2.72 | 0.12 | 2.82 | 0.07 |
| | 16:0 9,16-dihydroxy | 0.00 | 0.00 | 0.00 | 0.00 | 9.64 | 1.25 | 7.41 | 0.47 | 5.77 | 0.08 | 6.15 | 0.18 | 7.98 | 0.62 | 9.43 | 0.58 | 10.68 | 0.88 | 11.14 | 0.32 |
| | 18:0 18-hydroxy | 0.00 | 0.00 | 0.00 | 0.00 | 0.00 | 0.00 | 2.78 | 0.15 | 2.30 | 0.09 | 1.81 | 0.13 | 1.11 | 0.03 | 0.66 | 0.02 | 0.61 | 0.07 | 0.64 | 0.02 |
| | 18:2 9-hydroxy | 19.10 | 6.83 | 14.81 | 2.45 | 7.24 | 0.80 | 3.27 | 1.06 | 1.08 | 0.19 | 0.88 | 0.14 | 0.86 | 0.09 | 1.19 | 0.37 | 1.55 | 0.41 | 1.91 | 0.53 |
| | 18:1 18-hydroxy | 0.00 | 0.00 | 0.00 | 0.00 | 17.74 | 6.09 | 32.77 | 2.51 | 37.16 | 4.46 | 26.06 | 2.95 | 26.39 | 3.43 | 22.24 | 2.42 | 19.48 | 0.63 | 17.69 | 0.99 |
| | 18:0 9,10, 18-trihydroxy | 0.00 | 0.00 | 0.00 | 0.00 | 0.00 | 0.00 | 0.00 | 0.00 | 0.38 | 0.03 | 0.52 | 0.09 | 0.62 | 0.10 | 0.80 | 0.03 | 0.92 | 0.23 | 1.02 | 0.24 |
| | 18:1 10 (9),18-dihydroxy | 0.00 | 0.00 | 0.00 | 0.00 | 4.88 | 0.57 | 11.20 | 2.09 | 14.14 | 1.25 | 20.21 | 1.28 | 18.27 | 1.76 | 19.02 | 2.32 | 18.70 | 2.04 | 19.26 | 1.77 |
| | 18:0 9-epoxy-18-hydroxy | 0.00 | 0.00 | 0.00 | 0.00 | 2.34 | 0.20 | 3.69 | 0.44 | 7.42 | 0.48 | 11.71 | 0.67 | 11.84 | 0.44 | 11.21 | 0.32 | 11.15 | 0.89 | 10.74 | 1.39 |
| **α,ω-Dicarboxylic acids** | | 0.00 | 0.00 | 0.00 | 0.00 | 0.00 | 0.00 | 0.00 | 0.00 | 0.00 | 0.00 | 0.00 | 0.00 | 0.00 | 0.00 | 0.00 | 0.00 | 0.00 | 0.00 | 0.00 | 0.00 |
| | 16:0 | 34.67 | 4.43 | 43.54 | 2.53 | 23.14 | 0.34 | 8.48 | 0.43 | 4.38 | 0.02 | 3.05 | 0.10 | 2.08 | 0.04 | 1.47 | 0.10 | 1.42 | 0.03 | 1.58 | 0.12 |
| | 17:0 8-hydroxy | 0.00 | 0.00 | 0.00 | 0.00 | 0.00 | 0.00 | 0.00 | 0.00 | 0.00 | 0.00 | 0.46 | 0.09 | 0.35 | 0.00 | 0.33 | 0.02 | 0.29 | 0.02 | 0.32 | 0.01 |
| | 18:1 9-hydroxy | 0.00 | 0.00 | 0.00 | 0.00 | 7.39 | 0.43 | 13.36 | 1.12 | 15.53 | 0.32 | 18.40 | 0.54 | 21.73 | 2.17 | 26.30 | 1.72 | 27.66 | 1.83 | 27.65 | 2.12 |
| | 18:3 | 0.00 | 0.00 | 0.00 | 0.00 | 0.00 | 0.00 | 1.56 | 0.07 | 1.80 | 0.24 | 1.27 | 0.15 | 1.29 | 0.17 | 1.06 | 0.11 | 0.92 | 0.05 | 0.88 | 0.05 |
